# Maximizing Nature-based Solutions using Artificial Intelligence to align global biodiversity, climate, and water targets

**DOI:** 10.1101/2025.01.06.631540

**Authors:** Camilo Alejo, Amy Luers, Andréa Ventimiglia, María-Isabel Arce-Plata, H. Damon Matthews

## Abstract

Nature-based Solutions (NbS) encompass a spectrum of conservation and restoration actions aimed at improving biodiversity, climate, and/or water outcomes. Considerable research exists that focuses either on conservation or restoration, or on one particular environmental outcome. Yet, there is a need to develop integrated frameworks that align multiple outcomes and enable comprehensive environmental and economic assessments. Here, we present an integrated framework leveraging an AI agent that interprets species’ habitat connectivity changes along with climate and water co-benefits to select optimal conservation and restoration priorities. We implement this framework through scenarios maximizing biodiversity protection with ecological integrity, carbon storage, and water co-benefits to achieve Canada’s 30×30 conservation and restoration targets. Our results suggest that prioritizing the protection of threatened biodiversity and irrecoverable carbon storage would optimally enhance existing Protected Areas outcomes and conserve 30% of lands by 2030. However, effectively restoring 30% of degraded land will require targeted actions in existing natural and transformed ecosystems. The assessment of current anthropogenic pressures suggests that conservation and restoration actions may enhance climate resilience for forestry in natural lands and potentially benefit agricultural production and public health in transformed lands. Moreover, the mining sector represents the largest growing pressure on both conservation and restoration priorities in the upcoming decade. Overall, this integrated framework reveals strategic conservation and restoration priorities to align environmental targets, while identifying opportunities to coordinate NbS interventions across sectors.

## Introduction

Global biodiversity loss is closely intertwined with climate change, water scarcity, and social inequality. Given these interconnections, many global targets emphasize the need for pathways that simultaneously address environmental and social challenges. Centered around biodiversity, the Kunming-Montreal Framework aims to restore 30% of degraded land and establish 30% of area-based conservation by 2030 (i.e., 30×30 targets), ensuring enhanced biodiversity, ecosystem integrity, and related co-benefits (CBD, 2022). Since the establishment of the Paris Agreement (UNFCCC, 2015), reducing land use emissions while conserving and restoring ecosystem carbon sinks have become relevant components of Nationally Determined Contributions (NDCs) to reach net-zero emissions by 2050. Similarly, the Sustainable Development Goals (SDGs) set targets to protect and restore ecosystems, including water-related ecosystems such as forests, wetlands, rivers, and lakes (United Nations, 2015).

Nature-based Solutions (NbS) offer a broad and strategic spectrum of actions in ecosystems (Cohen-Shacham et al. 2016) to reach global environmental targets. These actions encompass protecting intact and natural ecosystems, sustainable management of semi-natural ecosystems, and actively restoring ecosystems that have been heavily modified by agriculture, urbanization, and other industrial activities (Griscom et al. 2017; Seddon et al. 2020). NbS therefore provide pathways for countries to reach the 30×30 conservation and restoration targets (CBD 2022) in synergy with climate change mitigation (Matthews et al. 2022), water management (Ferreira et al. 2023), and Sustainable Development Goals (Gómez Martín et al. 2020). Simultaneously aligning these environmental outcomes require analytical tools to identify optimal conservation and restoration opportunities across diverse landscapes.

Over the past decade, analytical tools for NbS monitoring and prioritization have become increasingly available. State-of-the-art monitoring tools typically leverage field observations, remote sensing, and machine learning to predict spatial-temporal patterns of biodiversity (Chollet Ramampiandra et al., 2023), carbon stocks (Sothe et al., 2022; Walker et al., 2022), and water quality (Chen et al., 2022). In parallel with this increased monitoring capacity, other approaches identify optimal NbS priorities for conservation and restoration actions. For instance, conservation priority frameworks have weighed the importance of biodiversity aspects such as irreplaceability (e.g., species and habitats rarity) and vulnerability through a reactive (e.g., threatened species, habitat loss) or proactive (low vulnerability and low cost) approach (Brooks et al., 2006; Funk & Fa, 2010). From an ecosystem services perspective, Mitchell et al. (2021) prioritized conservation co-benefits emerging from carbon storage, water supply, and tourism.

Others have aimed to prioritize conservation targets by aligning biodiversity outcomes with climate suitability (Eckert et al., 2023; Saunders et al., 2023; Stralberg et al., 2020), carbon storage (Soto-Navarro et al., 2020), or multiple co-benefits simultaneously (Jung et al., 2021; Neugarten et al., 2024; O’connor et al., 2021). These approaches rely on current species’ area of habitat, typically overlooking changes in habitats and connectivity that could more reliably distinguish conservation actions from alternative NbS options.

The identification of optimal NbS has not been restricted to conservation targets. For instance, biodiversity and carbon outcomes have been ranked and prioritized on lands that have been converted to pastures and croplands (Strassburg et al., 2020). Vettorazzi & Valente (2016) proposed land suitability assessments for prioritizing restoration actions that could improve water supply. A common approach between conservation (Chauvenet et al., 2020) and restoration (Mappin et al., 2019) has resulted from prioritizing vulnerability (e.g., habitat conversion, human footprint) on global ecoregions. Moreover, the potential of Artificial Intelligence (AI) to find optimal solutions in simulated and real-world environments has recently begun to be explored in either conservation (Silvestro et al., 2022) or restoration (Currie et al., 2023; Silvestro et al., 2025) priority frameworks. These advancements in NbS prioritization represent an opportunity to develop integrated frameworks for conservation and restoration that leverage AI to maximize biodiversity and related co-benefits outcomes.

The complexity of optimizing biodiversity and co-benefit outcomes for conservation and restoration targets is further challenged by the interplay of economic priorities. For example, some priority frameworks highlight the restoration potential of converted lands for pasture and agriculture to reach environmental targets (Currie et al., 2023; Strassburg et al., 2020). However, realizing this potential may be unfeasible due to economic barriers (Cook-Patton et al., 2021) and may require considering other degraded lands to reach restoration targets. Similarly, prioritizing natural resource extraction will limit governments’ pathways to achieve environmental targets while diminishing current biodiversity and related co-benefits outcomes (Carr et al., 2021; Lawley et al., 2022; Rehbein et al., 2020). Tackling this complexity requires exploring diverse conservation and restoration scenarios that enable comprehensive assessments of environmental and economic implications.

Here, we present an integrated framework to prioritize and assess the environmental and economic implications of potential conservation and restoration areas. Our framework employs Reinforcement Learning, where an agent is trained to learn from system states (i.e., exploration) and cost-effectively select areas (i.e., exploitation), maximizing pre-determined rewards. In our case, the agent relies on pre-determined assessments of species’ habitat and connectivity changes, along with climate and water co-benefits, to perform a cost-effective selection that maximizes these environmental outcomes. We test this framework in Canada under a series of scenarios to reach 30×30 conservation and restoration targets that maximize the occurrence of threatened species in synergy with co-benefits from ecological integrity, irrecoverable carbon storage, and water surface stability. Overall, our findings suggest that the optimal scenario to conserve 30% of lands in Canada requires protecting threatened biodiversity and carbon stocks. Restoring 30% of degraded lands is more challenging, requiring a combination of scenarios across disturbed natural and transformed ecosystems. An assessment of anthropogenic pressures across conservation and restoration scenarios suggests intragovernmental and inter-sectoral coordination is needed. Taken together, our results demonstrate that a framework for conservation and restoration that integrates AI, multiple co-benefits, and scenario analysis can reveal synergistic and optimal NbS to achieve global environmental targets.

## Materials and methods

Our study aims to leverage geospatial analysis and Reinforcement Learning to prioritize NbS locations and reach Canada’s 30×30 targets while identifying synergies with other co-benefits. To this end, we explored two sets of NbS priority scenarios that maximize the occurrence of threatened vertebrate and plant species within selected land areas (Figure 1). The first set of scenarios focuses on conservation priorities to protect threatened biodiversity habitats and reach 30×30 conservation targets (i.e., conserve 30% of land by 2030) in areas with high ecological integrity (Conservation Scenario 1), high and irrecoverable carbon stocks (Conservation Scenario 2), and stable surface water (Conservation Scenario 3). The other set of priorities is based on recovering threatened biodiversity habitats to reach 30×30 restoration targets (i.e., restore 30% of degraded land by 2030) in areas that have lost ecological integrity (Restoration Scenario 1), carbon stocks (Restoration Scenario 2), and surface water stability (Restoration Scenario 3). Based on the resulting priority scenarios, we analyze the spatial patterns of conservation and restoration priorities across ecological and administrative boundaries. We then compare these patterns to current and future natural resource extraction activities, agriculture, and urban development.

**Figure 1.**
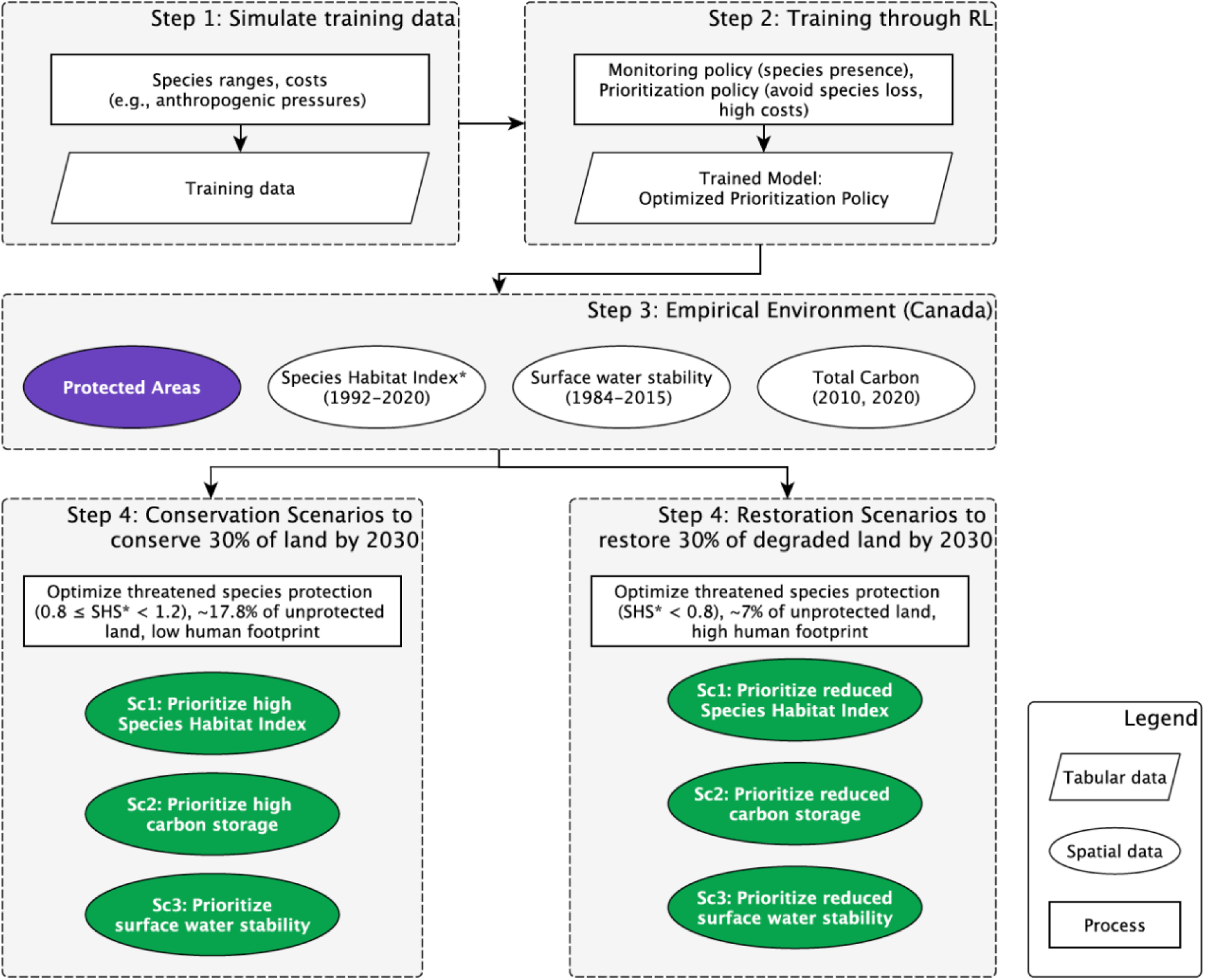
Flowchart summarizing the steps to identify Conservation and Restoration priorities to reach 30×30 targets aligned with co-benefits, using Reinforcement Learning. *The Species Habitat Index averages Species Habitat Scores (i.e., SHS). These scores account for species’ changes in area of habitat and connectivity and allow for determining suitable areas for conservation or restoration.

### Study Area

We use Canada as a case study, which has the second-largest national land mass with more than 9.9 million square kilometers of surface area (Statistics Canada, 2011). It is estimated that Canada is home to around 80,000 species, including mammals, birds, fishes, amphibians, reptiles, plants, and invertebrates (Canadian Endangered Species Conservation Council, 2016). These species are found across diverse ecozones that contain approximately ∼25% of global wetlands, boreal forests, and temperate forests (Statistics Canada, 2011). Summing aboveground biomass, belowground biomass, and soil organic carbon, Canada’s carbon stocks are estimated at 33.7 PgC, representing 13.7% of global carbon storage, surpassed only by Brazil (18.2%) and Russia (20.9%) (Walker et al., 2022). Moreover, Canada has a surface water area of 1.1 million square kilometers, accounting for 26% of the global surface water distribution (Pekel et al., 2016). Given these ecological and geographic features, Canada presents significant potential to implement NbS initiatives aimed at achieving combined biodiversity, carbon storage, and surface water outcomes.

Canada is also considered to have one of the most extensive areas of global intact ecosystems (Watson et al., 2018; Williams et al., 2020). The current network of Protected Areas covers 12.7% of Canada’s terrestrial land (Figure S1), partially securing these ecosystems’ biodiversity, carbon stocks, and surface water (Environment and Climate Change Canada, 2023). However, within Canada’s boundaries, almost 25% of land area is experiencing human pressures (Hirsh-Pearson et al., 2022), and 10% of vertebrate and plant species registered by the International Union for Conservation of Nature red list are classified under some category of extinction risk (IUCN, 2023). Additionally, significant portions of Canada’s carbon stocks are at risk of being lost and not recovering by 2050 due to land use and climate change, particularly in ecozones along the Pacific (e.g., Pacific Maritimes), Southern Arctic, and Hudson Plains (Noon et al., 2022). Surface water faces similar risks, which have resulted in transitions from permanent water cover to seasonal or complete loss (Pekel et al., 2016).

### Data processing

To determine NbS priority scenarios of conservation and restoration through Reinforcement Learning, we used a range of datasets on the spatial patterns of biodiversity, carbon stocks, and surface water occurrence in Canada (Table S1). We used the IUCN species range polygons to establish the approximate spatial distribution of mammals, amphibians, reptiles, and plant species in terrestrial ecosystems (IUCN, 2023). Bird species range polygons were obtained from BirdLife International (BirdLife International & Handbook of the Birds of the World, 2022). Using IUCN’s and BirdLife’s range polygons and associating species’ habitat preferences (Brooks et al., 2019) with ESA-CCI land cover (Defourny et al., 2023), and elevation maps (Natural Resources Canada, 2011), we refined species ranges to create Area of Habitat Maps for 1992 and 2020 with the R package aoh (Hanson, 2025). We opted for the ESA-CCI maps because they provided the longest consistent land cover monitoring product. For further processing, the Area of Habitat Maps were resampled to a 1km resolution. Following the IUCN Red List categories: Near Threatened, Vulnerable, Endangered, and Critically Endangered, we also established which species were experiencing some risk of extinction. According to these categories, birds (n = 48), and mammals (n = 11) represented 88% of threatened species (n = 59), while the remaining 12% corresponded to plants (n = 6), reptiles (n = 1), and amphibians (n = 1) (Figure S2).

Based on the Area of Habitat of threatened and non-threatened species, we estimated a Species Habitat Index as a proxy of ecological integrity (Figure S3). The Species Habitat Index measures changes in the quality and connectivity of habitats for species of interest relative to a baseline (Arce-Plata et al., 2025; Jetz et al., 2022; Jetz & GEO BON Secretariat, 2002). In our case, we assessed habitat quality by measuring changes in the Area of Habitat for each species between 1992 and 2020 within 10×10 km planning units grids. To assess habitat connectivity, we measured the average changes in the distance to species’ habitat edge in the same temporal and spatial scale with the R package landscapemetrics (Maximillian & Hesselbarth, 2021). The resulting habitat quality and connectivity scores were averaged to obtain a Species Habitat Score (SHS), suggesting the decrease, stability, or increase in suitable area for a single species. Among threatened species, the SHS allowed us to determine suitable areas for conservation (0.8 ≤ SHS < 1.2) and restoration (SHS < 0.8) for single species.

Subsequently, the SHS were averaged for threatened and non-threatened species to estimate the Species Habitat Index, providing an estimate of ecological integrity in 2020 relative to 1992 within 10×10 km planning units across Canada (Figure S4).

Total carbon annual stocks and changes were estimated using biomass, soil organic carbon, and land cover maps (Table S1). Data sources included a 2010 global harmonized biomass carbon map (∼300m resolution) from Spawn et al. (2020) and soil organic carbon data (∼250m resolution) from Sothe et al. (2022). The latter includes estimates from 2000-2019 averages at 30 cm and 100 cm of depth in Canada. To determine total carbon changes, ESA CCI annual land cover maps (Defourny et al., 2023) were converted to 9 IPCC classes following Hu et al. (2021). The temporal timeframe of carbon data allowed us to use class transitions between 2010 and 2020 to estimate biomass and soil organic carbon changes separately (Table S2).

We calculated total carbon stocks losses, using the following approach. Vegetation-to-non vegetation (e.g., forest-urban) transitions resulted in a complete biomass loss. For vegetation-to-vegetation transitions, partial carbon losses were calculated as the difference between initial high-carbon vegetation (e.g., forest) and the ecozone average of low-carbon vegetation (e.g., grassland). We assumed soil organic carbon remained stable in vegetation-only transitions (W. Li et al., 2018; Noon et al., 2022). Using peatland maps (Xu et al., 2018), the transitions from high-carbon vegetation classes to agriculture or urban or bare land involved a carbon loss function in non-peat soils at 30 cm of depth and complete carbon loss in peat soils at 100 cm of depth (Noon et al., 2022; Poeplau et al., 2011).

Similarly, total carbon gains involved region-based estimates. Biomass gains were calculated using sequestration rates for boreal, temperate lowland, temperate mountain (Noon et al., 2022), and tundra regions (Rocha & Shaver, 2011). Soil organic carbon gains (e.g., agriculture to shrubland) were estimated using recovery functions in non-peat soils at 30 cm depth for temperate, boreal (Poeplau et al., 2011), and tundra regions (Loranty et al., 2014). The recovery functions in boreal and temperate peatlands corresponded to soil organic carbon estimates at 100 cm depth (Poeplau et al., 2011).

Wetland-forest transitions were excluded due to potential classification errors and temporal transitions (Amani et al., 2021; N. Li et al., 2023). These changes in biomass and soil organic carbon in 2010 and 2020 were combined to create annual total carbon maps (Figure S5) and to estimate annual stock changes (Figure S6) at a 1km resolution. To match the spatial scale of planning units, both annual and change estimates of total carbon were resampled to a 10 km resolution.

In addition to the spatial patterns of threatened species, ecological integrity, and total carbon stocks, we quantified water surface changes (Figure S7). We used the Landsat-derived estimates of changes in global surface water from 1984 to 2015 (Pekel et al., 2016). This dataset exhibits the spatial and temporal changes of any stretch of open, fresh, and salt water, larger than 30m by 30m. Specifically, we focused on water occurrence change intensity that results from comparing the frequency of monthly water detection between two periods: 1984-1999 and 2000-2015. Thus, the water occurrence change intensity provides a measure of gains, losses, and persistence. After masking other land covers, we estimated the mean water occurrence change intensity in sub-basins nested in hydrological basins across Canada. Specifically, we used HydroBASINS level 5 sub-basin boundaries (Lehner & Grill, 2013), which represent an intermediate water-shed subdivision and may exhibit regional impacts of domestic, industrial agricultural water use (Schlattmann et al., 2022). Overall, capturing these long-term changes in water occurrence may reveal the impacts of land use and climate change while highlighting adaptation actions affecting one of the most accessible sources of water for human populations (Pekel et al., 2016). Surface water changes were estimated in Google Earth Engine, and the geospatial processing of biodiversity, ecological integrity and total carbon stocks data were executed with the R package Terra (Hijmans et al., 2024).

### Reinforcement learning to determine conservation and restoration priorities

We used the Reinforcement Learning algorithm CAPTAIN (Silvestro et al., 2022) to identify conservation and restoration priorities to reach 30×30 targets in Canada. The process of identifying priorities with this algorithm can be summarized in four steps: (1) simulate hypothetical spatial training data; (2) train an optimized model of prioritization through Reinforcement Learning; (3) build an empirical environment using observed real-world data; and (4) apply the optimized model within the empirical environment to identify priority conservation and restoration areas (Figure 1).

In the first step, hypothetical natural environments are simulated with a resolution, spatial extent, and several hypothetical species that resemble the empirical environment. Each hypothetical species displays unique spatial distributions, demographic traits, dispersal capacity, and sensitivity to climate and anthropogenic disturbances. Based on the hypothetical environments, we implemented a monitoring policy, a budget (target), and a priority policy to be used for model training. In our case, the monitoring policy gathers information in each planning unit about species (including their presence) and a cost, which could be interpreted as a level of anthropogenic pressure. The budget defines the maximum fraction of planning units that can be prioritized and is a function of the anthropogenic costs. Here, the budget was arbitrarily defined as 10%, that is, a tenth of the total cost of all planning units in the simulated natural environment. Constrained by the budget, the prioritization policy was set to reward the selection of planning units that maximize the occurrence of species and guarantee at least 10% of the spatial distribution of all species while avoiding high costs (e.g., high anthropogenic pressures).

In the second step (Figure 1), an optimization model was trained to prioritize planning units (i.e., spatial 10×10 km grids) using Reinforcement Learning according to the predetermined monitoring policy, budget, and prioritization policy. To this end, CAPTAIN optimizes the parameters of a neural network to cost-effectively maximize the rewards resulting from prioritizing planning units. As explained in detail by Silvestro et al.(2022), this training process relies on a two-phase parallelized evolution strategy that repeats in several epochs. During the first phase, changes in policy parameters are randomly perturbed and tested in a simulated change in the environment (i.e., adding or removing prioritized planning units), resulting in new priority actions and a reward. After simulating these changes in different iterations that occur in parallel, the second phase aggregates the results of these iterations to identify improvements in the rewards and update the neural network parameters. The two-phase process repeats across different epochs (here, 100), and the rewards from a current epoch are compared with the weighted average results of previous epochs in a process defined as an advantage function.

Thus, based on the simulated natural environments, we produced a trained optimization model capable of cost-effectively identifying priority planning units that maximize the occurrence of species.

In the third step, we integrated observed spatial datasets for Canada as an empirical environment. This environment was created by dividing the country’s land surface into planning units of 10×10 km that included information regarding suitable areas of threatened species, the occurrence of Protected Areas, ecological integrity, total carbon stocks, total carbon stocks changes, and water occurrence change intensity. Using the Canadian Protected and Conserved Areas database (Environment and Climate Change Canada, 2023), we excluded planning units with a Protected Area surface equal to or higher than 50 km2, resulting in 12.25% of the area assumed to be already protected and therefore excluded from priority scenarios. Estimates regarding ecological integrity, carbon stocks and water occurrence change intensity across planning units were normalized to create a scale of costs for conservation and restoration priority scenarios.

In the final step, the priority scenarios for conservation and restoration aimed to reach 30×30 targets that would maximize the occurrence of threatened biodiversity, guaranteeing at least 10% of threatened species’ ranges while avoiding specific costs. We aimed to reach 30×30 targets for the conservation scenarios by identifying at least ∼16600 priority planning units (∼17.75% of the total area) where threatened species exhibited stable or moderate changes in the Species Habitat Scores (0.8 ≤ SHS < 1.2) (Figure S3). In Conservation Scenario 1, the normalized ecological integrity values were recalculated so that the highest value (∼ 1), and therefore the highest cost, corresponded to the lowest ecological integrity. Similarly, the Conservation Scenario 2 was based on a cost scale that rewarded the selection of low cost-high total carbon. To prioritize carbon stocks that, if lost, would not recover by 2050 (Noon et al., 2022), the original cost of planning units found in areas with irrecoverable carbon stocks was further reduced to a tenth of the original value. Using a cost scale displaying the largest values for the most intense water occurrence changes, the Conservation Scenario 3 rewarded the prioritization of planning units displaying low cost-low water surface change. To further reward water surface stability, we reduced to a tenth the original value of planning units that experienced an absolute water change below 20%. In all scenarios, the cost of planning units in areas with moderate to high human footprint (<5) (Hirsh-Pearson et al., 2022) was increased to the maximum cost value (1).

Consequently, the conservation scenarios prioritized ecological integrity, climate change mitigation potential, and surface water stability, suggesting proactive NbS with low anthropogenic pressures.

The restoration scenarios focused on areas that exhibited a reduction in threatened species Habitat Scores (SHS < 0.8) between 1992 and 2020 (Figure S3). The three scenarios aimed to identify priorities areas to restore 30% of degraded land. We assumed that all lands exhibiting some human pressure according to Canada’s human footprint (Hirsh-Pearson et al., 2022) were restorable, which corresponded to ∼7% of the total land area (∼6600 planning units). While the restoration scenarios had the same themes as the conservation scenarios, we used a different scale of costs and rewards for the prioritization process. Restoration Scenario 1 and Scenario 3 were based on scales of costs that rewarded the prioritization of planning units with low ecological integrity and high-water occurrence change intensity, respectively. Using a normalized scale where values closer to zero represented the largest carbon stock losses, the Restoration Scenario 2 rewarded the prioritization of areas that have experienced carbon stocks losses over the period 2010-2020. Similar to the Conservation Scenario 2, the cost of carbon stocks losses in planning units with irrecoverable carbon stocks (i.e., irrecoverable by 2050 via natural processes) was reduced to a tenth of the original value. By prioritizing reduced ecological integrity, carbon stock losses, and surface water instability, the restoration scenarios produced priority areas for reactive NbS requiring active management and adaptation actions under high anthropogenic pressures. Each conservation and restoration scenario underwent ten replicates, generating selection probabilities that guided the removal of excess planning units beyond the area targets.

### Conservation and Restoration priorities relative to current and future anthropogenic pressures

Finally, we compared the spatial distribution of resulting conservation and restoration priorities with current and future anthropogenic pressures. Current anthropogenic pressures included agriculture, grasslands, and urban development obtained from the 2020 Annual Crop Inventory (Agriculture and Agri-Food Canada, 2020). We also considered anthropogenic pressures from mining, oil and gas (Natural Resources Canada, 2020), and dams and associated reservoirs (Lehner et al., 2024; Lehner & Grill, 2013). Forestry between 1985 and 2020 derived from Landsat time series that distinguish forest disturbances from harvest and fire (Hermosilla et al., 2016). In addition to current anthropogenic pressures, the Major Projects Inventory provided the location of forestry, mining, and energy sector projects currently under construction or planned within the next ten years (Natural Resources Canada, 2024). This inventory included high-impact projects with a minimum investment between 20 (forestry and electricity) and 50 (mining and energy) million CAD. Based on this data, we identified the nearest distance of each conservation and restoration priority to current and future anthropogenic pressures in multiple sectors. By analyzing this spatial distribution, we aimed to identify current and potential land conflicts and relevant economic sectors for conservation and restoration actions.

## Results

### Scenarios to conserve 30% of land by aligning biodiversity, climate, and water outcomes

We defined conservation priority scenarios (Figure 1) based on the spatial distribution of existing Protected Areas (Figure S1), threatened species (Figure S2), ecological integrity (Figure S4), carbon storage (Figure S5), and surface water stability (Figure S7). The conservation scenarios identified an additional ∼17.75% of Canadian land areas to be protected. These scenarios achieve Canada’s 30×30 targets by maximizing the protection of threatened biodiversity and guaranteeing at least 10% of each threatened species range, while also prioritizing co-benefits of ecological integrity (Conservation Scenario 1), irrecoverable carbon stocks (Conservation Scenario 2), and surface water stability (Conservation Scenario 3) (Figure 2).

**Figure 2.**
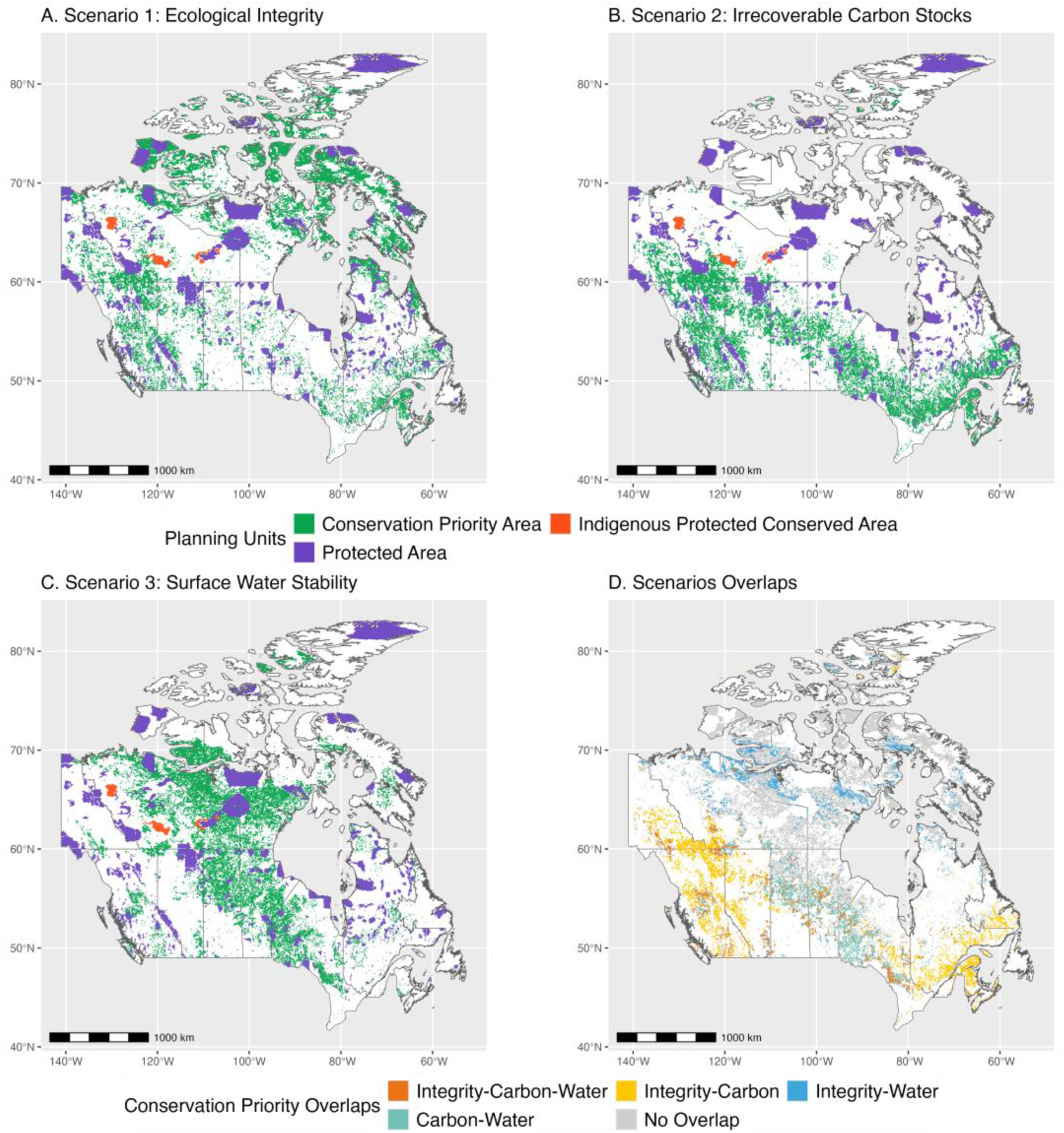
The spatial distribution of Conservation Priority Scenarios to conserve 30% by 2030 obtained through Reinforcement Learning. All Scenarios assume a baseline extent of Protected Areas of 12.25% and a 17.75% remaining area budget to maximize the occurrence of threatened species with co-benefits emerging from Ecological Integrity (A), Irrecoverable Carbon Stocks (B), Water Surface Stability (C). Priority Areas had a 10km resolution. The Scenarios Overlaps (D) suggest areas that simultaneously maximize the occurrence of threatened species with multiple co-benefits.

When ecological integrity was prioritized (Conservation Scenario 1; Figure 2), more than 30% of conservation priorities were concentrated in the Southern and Northern Arctic (Figure S8) (i.e., Quebec, Nunavut, Northwest Territories) and 20% in Boreal ecozones (i.e., Ontario, Quebec, Alberta, British Columbia) (Figure S9). Relative to existing Protected Areas, this scenario would result in an expected average increase of ecological integrity from 0.64 to 0.84 on a normalized scale (Figures 3, S10). Similarly, prioritizing ecological integrity would increase average carbon storage (279.8 to 350 tC/ha). However, this scenario would reduce the average number of threatened species protected (7.73 to 5.6 species/100 km^2^), species richness (83.19 to 70.66 species/100 km2), and absolute water surface stability (9.87 to 10.47 % in 2000-2015 vs 1984-1999) (Figure 3).

**Figure 3.**
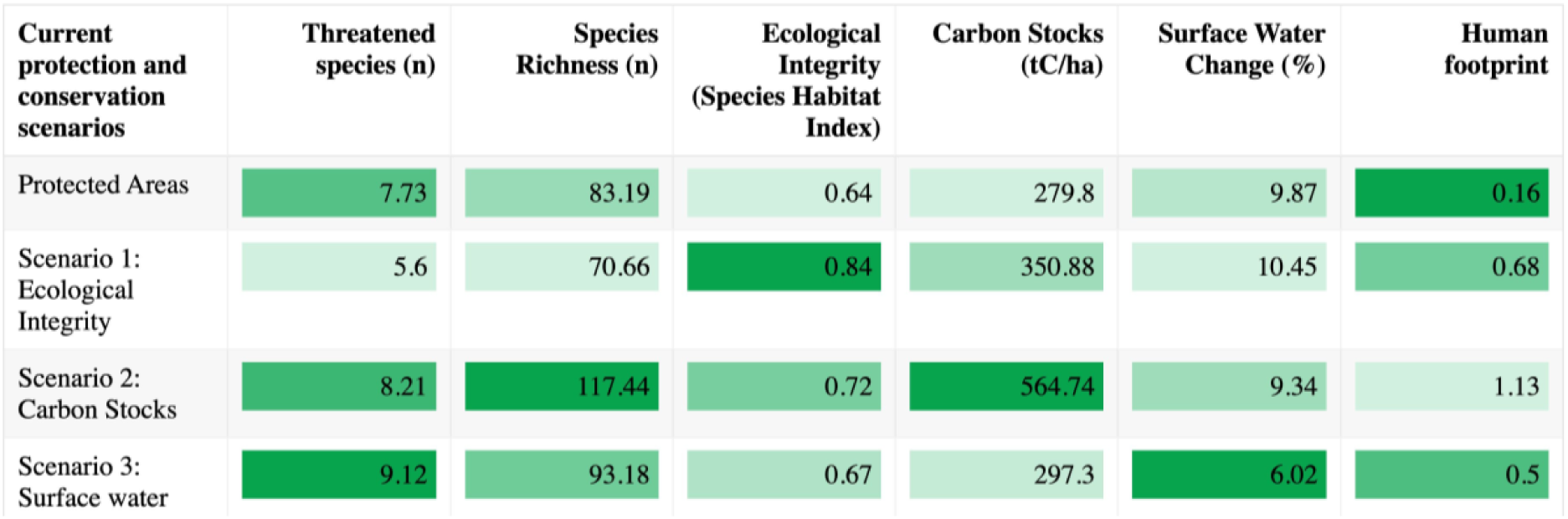
The average outcomes of existing Protected Areas and Conservation Priority Scenarios in threatened species, species richness, ecological integrity, carbon stocks, and absolute surface water change. The conservation scenarios aim to maximize the occurrence of threatened species with co-benefits emerging from Ecological Integrity, Irrecoverable Carbon Stocks, and Water Surface Stability.

Conservation Scenario 2 (prioritizing irrecoverable carbon stocks; Figure 2) led to conservation priorities that extend across the Atlantic Maritimes (4.6%) to the forest-dominated Boreal ecozones (∼ 60%) and the western Montane Cordillera (∼11%) (Figure S8) throughout eastern, central, and western provinces (Figure S9), respectively. In contrast to Scenario 1, aligning biodiversity and climate change mitigation targets led to slightly lower ecological integrity (0.72) and an increase in the human footprint (i.e., anthropogenic pressure score 1.12/55). Compared to Protected Areas, this co-benefit consistently improved as the number of protected threatened species (8.21 species/100km^2^), species richness (117.44 species/100km^2^), water surface change (9.3%), and carbon stocks (564.74 tC/ha) (Figures 3, S10).

Conservation Scenario 3, which prioritized surface water stability in hydrological sub-basins (Figure 2), had a dispersed geographic distribution. A significant portion of priorities overlapped with Conservation Scenarios 1 and 2, especially in the Boreal ecozones and the Southern Arctic. Other priorities were particularly aggregated in the western Taiga Shield (Ontario, Manitoba, Northwest Territories) and Taiga Plains (British Columbia) (Figures S8, S9), known to be partially dominated by peatlands. Compared to existing Protected Areas and other scenarios, prioritizing surface water stability would increase the number of threatened species protected (9.12 species/100km2) and surface water stability (6.02%). This prioritization scenario would also improve species richness (93.18 species/100km^2^), ecological integrity (0.67), and carbon storage (197.3 tC/ha) outcomes beyond what existing Protected Areas achieve, though not as effectively as other scenarios (Figures 3, S10).

After overlapping the results from all three conservation scenarios (Figure 2), we were able to identify the areas of land that were prioritized in two or more of the individual scenarios. Of the target 17.75% of land area prioritized by the three scenarios, 12.5% appeared in two scenarios and 2.7% in all three scenarios, suggesting significant potential for achieving multiple co-benefits. These overlaps confirm the importance of Boreal (Churchill River, Saskatchewan, Manitoba; Lac Temiscamingue Lowland, Ontario; Abitibi Plains, Ontario and Quebec), Montane Cordillera (e.g., Omineca Mountains, Fraser Plateau, British Columbia) and Taiga (e.g., Northern Alberta Uplands) ecozones to achieve substantial outcomes for threatened biodiversity and other co-benefits. Furthermore, the overlapping conservation priorities also highlight carbon storage as a unifying co-benefit that aligns with other co-benefits and would strategically connect existing Protected Areas nationally.

The conservation scenarios overlap also reveal a reduced portion of strategic priorities that maximize environmental outcomes in limited-extent ecozones. For example, compared to existing Protected Areas, the overlapped priorities in the Atlantic Maritimes (0.68%, e.g., Southern New Brunswick Uplands), Prairies (0.30%, e.g., Alberta’s mixed grasslands), Hudson Plains (0.08%, e.g., James Bay Lowlands in Ontario and Québec), and Mixedwood Plains (0.03%, Quebec and Ontario) (Figures S7, S8) would simultaneously and significantly increase biodiversity outcomes, ecological integrity, carbon stocks, and surface water stability (Table S3). Overall, the Conservation Scenarios imply that aligning biodiversity outcomes with other co-benefits results in varied spatial patterns and some trade-offs. Nevertheless, maximizing the protection of threatened biodiversity and irrecoverable carbon stocks (Conservation Scenario 2) represents a comprehensive pathway to improve biodiversity, climate change mitigation, and water stability outcomes in existing Protected Areas.

### Scenarios to restore 30% of degraded land by aligning biodiversity, climate, and water outcomes

The restoration scenarios suggested priorities in 30% of degraded land (i.e., ∼7% total surface) that would maximize the occurrence of threatened biodiversity in areas experiencing reduced ecological integrity (Restoration Scenario 1), carbon stocks losses (Restoration Scenario 2), and surface water instability (Restoration Scenario 3) (Figure 4). Restoration Scenario 1 priorities were mostly concentrated in the Boreal ecozones (∼70%) (Figure S8), dominated by forest lands and wetlands. Among all Restoration Scenarios, prioritizing reduced ecological integrity was particularly effective for protecting threatened biodiversity (∼10.49 species/100 km^2^) and species richness in general (141.12 species/100 km^2^), and it was moderately effective for carbon storage (519.33 tC/ha) (Figure 5). When threatened biodiversity was prioritized with irrecoverable carbon stocks losses, a considerable part of the priorities occurred in Boreal (57%) and Montane Cordillera (25%) ecozones (Figure 4). Although Restoration Scenario 2 would result in slightly lower biodiversity outcomes than Restoration Scenario 1, prioritizing the recovery of carbon stock losses exhibited effective outcomes for ecological integrity (0.61), carbon stocks (597.41 tC/ha), and surface water stability (9.31%) (Figures 5, S11). The Restoration Scenario 3 displayed a significant number of priorities in the Prairies (56%) and Boreal ecozones (18%), displaying the highest outcomes for Ecological Integrity (0.86) and the lowest for carbon storage (279 tC/ha) among the three scenarios (Figure 5). Considering the moderate human footprint (8.2/55) when water surface instability was prioritized (Figure 5), the Ecological integrity values imply that in the past three decades, the species area of habitat and their connectivity have remained relatively stable.

**Figure 4.**
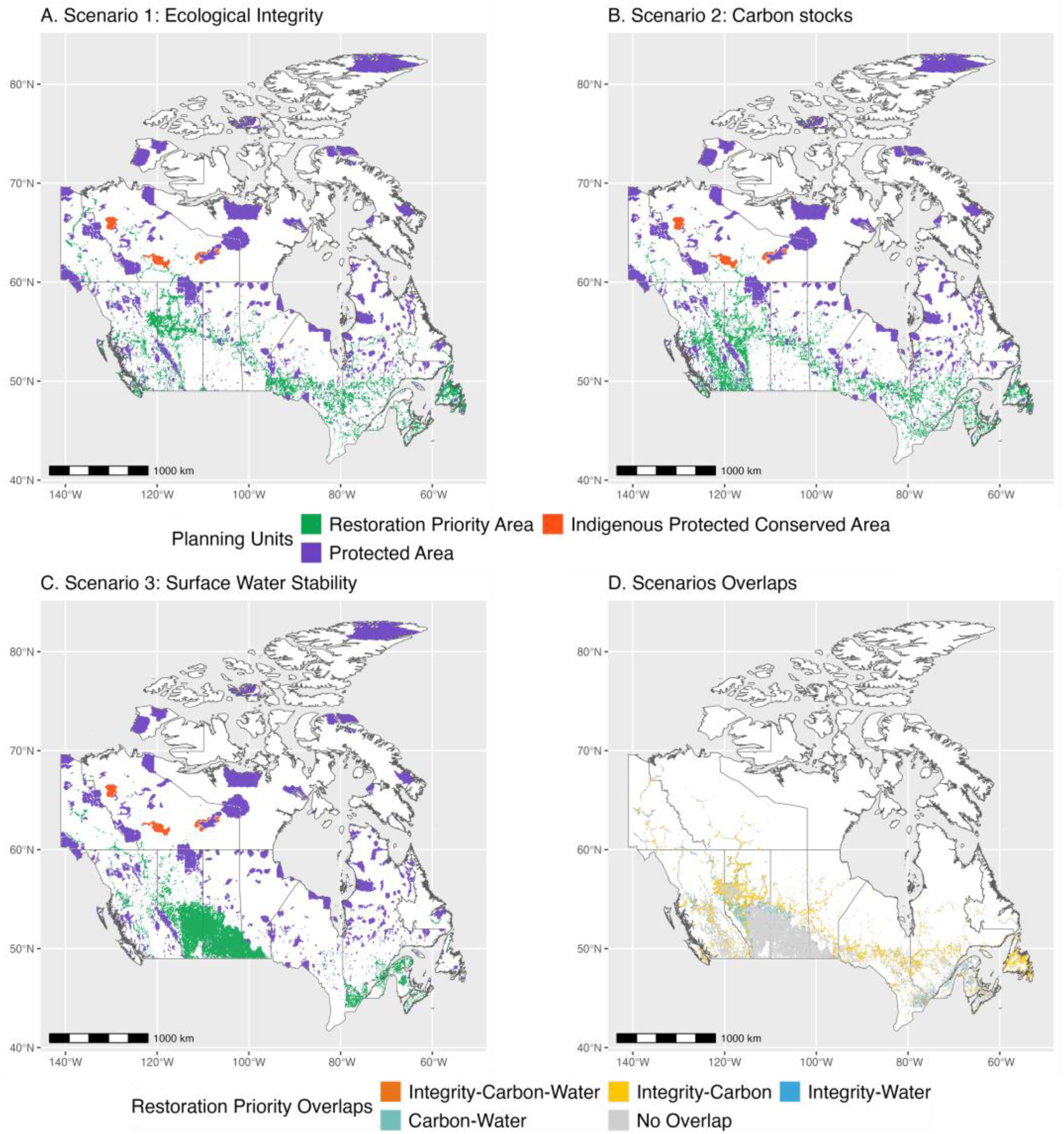
The spatial distribution of Restoration Priority Scenarios to restore 30% of degraded land by 2030 obtained through Reinforcement Learning. All Scenarios assume a baseline extent of degraded land of 23.5% and an area budget of ∼7% to maximize the occurrence of threatened species while prioritizing the loss of ecological integrity (A), carbon stocks (B), and water surface stability (C). Priority Areas had a 10km resolution.

**Figure 5.**
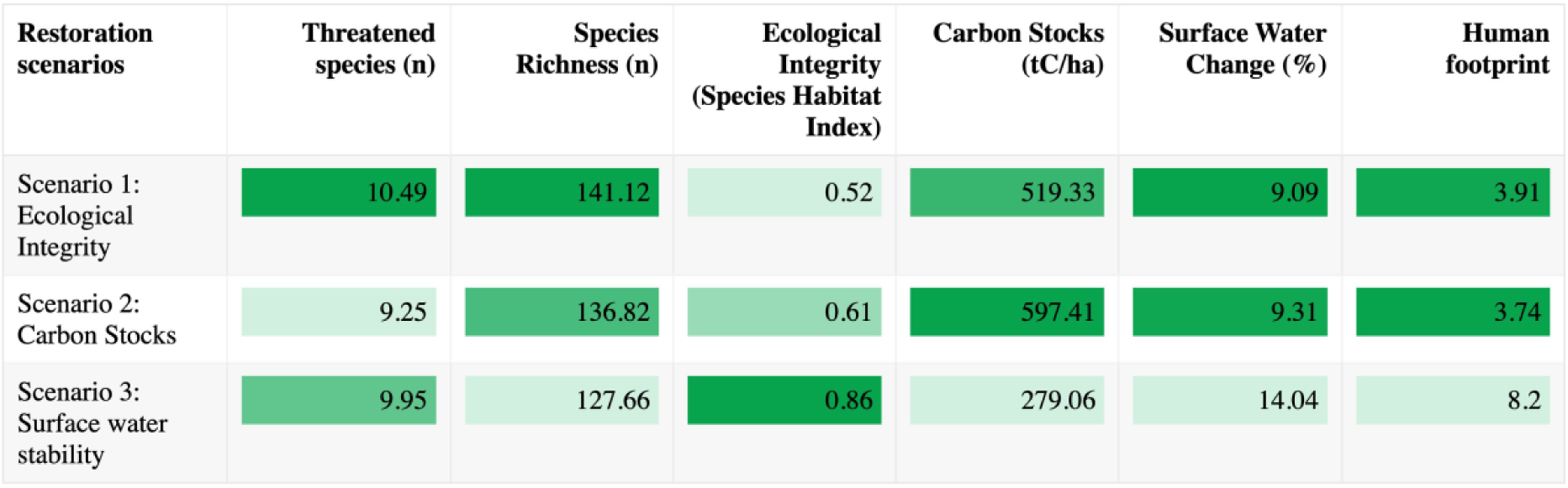
The average outcomes of Restoration Priority Scenarios in threatened species, species richness, ecological integrity, carbon stocks, and absolute surface water change. The restoration scenarios aim to maximize the occurrence of threatened species while prioritizing potential improvements in Ecological Integrity, Carbon Stocks, and Water Surface Stability.

The overlaps among Restoration Scenarios represented approximately two-thirds (4.67%) of the desired land target (7%) (Figure 4). Among these overlaps, only 0.16% simultaneously aligned potential improvements in ecological integrity, carbon storage, and water co-benefits. Notably, the restoration scenarios prioritizing ecological integrity and carbon storage displayed the largest overlap, representing nearly half of the potentially restorable land. The wide alignment of these two scenarios (Figure 4), mostly across Boreal ecozones (e.g., Boreal Plains in Northern Alberta, Boreal Shield in Ontario and Quebec) and the Montane Cordillera (e.g., Fraser Plateau, British Columbia), confirm their similar and high-quality environmental outcomes, relative to Protected Areas and conservation priorities (Figure 5). Additionally, a few of these overlapped priorities reveal strategic areas in under-represented ecozones. For example, some priorities in the Mixedwood plains (0.05%), Atlantic Maritimes (0.01%, e.g., Southwest Nova Scotia Uplands), and Pacific Maritimes (0.01%, e.g., Eastern Vancouver Island) displayed exceptionally high values for biodiversity, carbon and water outcomes (Table S4).

Overall, the Restoration Scenarios and their overlaps suggest two restoration pathways. According to Restoration Scenarios 1 and 2, one pathway implies coordinated actions for recovering threatened biodiversity, ecological integrity and carbon storage in ecosystems dominated by forests and wetlands with low to moderate human pressures. This pathway would result in short-term improvements in environmental outcomes.

Scenario 3, suggests a pathway focusing on recovering threatened biodiversity and surface water instability. The associated moderate to high human pressures in this pathway would result in mid to long-term improvements in environmental outcomes. In other words, both restoration pathways would potentially improve conditions for threatened biodiversity in tandem with one or multiple co-benefits in particular locations and time frames, suggesting diverse and context-dependent restoration actions.

### Conservation and restoration priorities are under varying pressures

To identify the current and future socio-economic implications of conservation and restoration scenarios, we assessed the spatial distribution of priorities in relation to different economic sectors. Regarding current implications, we quantified the distance of priority areas to agriculture, urban development, oil and gas, mining, forestry and dams (Figure 6). Forestry activities were within a 100 km radius of 59% of conservation priority areas in the scenario maximizing ecological integrity, and 69% of priority areas when maximizing surface water stability. The conservation scenario maximizing irrecoverable carbon storage exhibited the highest pressure from forestry (97% priority areas within a 100 km radius). This is an expected result, given that a significant number of conservation priority areas, especially those maximizing carbon storage, were in areas dominated by forest lands, such as the Boreal ecozones, Montane cordillera, and Atlantic Maritimes. Beyond forestry, the priority areas maximizing carbon storage experienced the highest pressure from hydroelectric dams (50%), followed by urban development, agriculture, mining (∼40%), and oil and gas (18%). These anthropogenic pressures are not spatially homogeneous and are particularly concentrated in the high human footprint Mixedwood plains (> 70%) (Table S6). Thus, forestry emerges as the most dominant current pressure on conservation priority areas, including those that would more effectively improve Protected Area outcomes.

**Figure 6.**
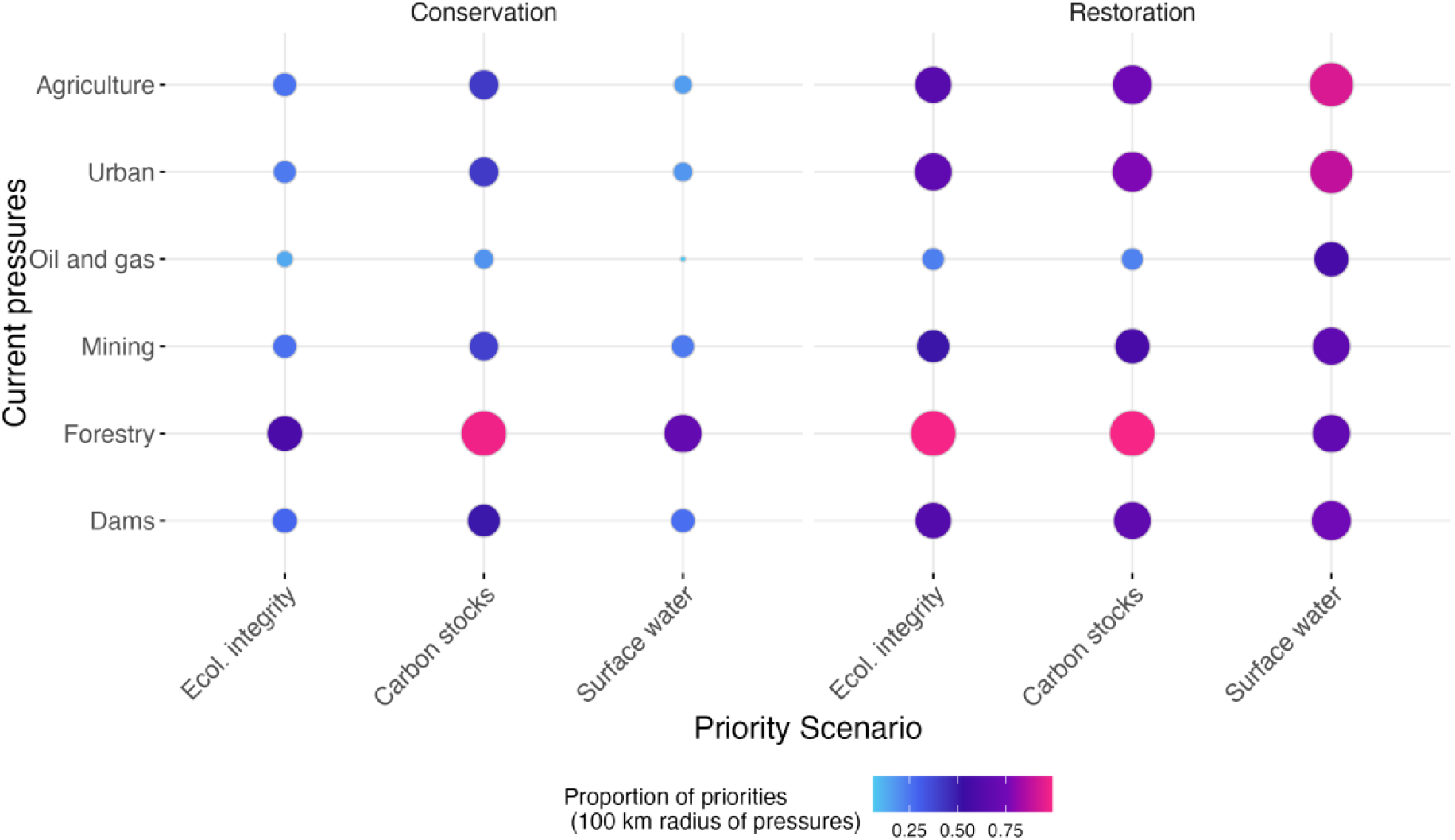
The proportion of conservation and restoration priority areas within a 100 km range of current anthropogenic pressures.

According to our scenarios, the restoration priority areas occur in non-intact areas with varying levels of human footprint, resulting in larger pressures compared to conservation priority areas (Figure 6). For example, 99% of restoration priority areas aiming at restoring habitats for threatened biodiversity, along with losses of ecological integrity and carbon storage, were within a 100 km radius of forestry activities. After forestry, the highest pressures on priority areas for restoring ecological integrity and carbon storage were agriculture (64% and 74%, respectively) and urban development (67% and 77%, respectively). The pressure of agriculture and urban development was more pronounced in priority areas aiming to restore habitats for threatened biodiversity and surface water stability (89-93%). These spatial patterns imply that restoration priority areas targeting surface water instability are strongly associated with areas dominated by agriculture, such as the Prairies and Mixedwood plains, or urban development in both the Mixedwood Plains and Atlantic Maritimes (Table S10).

Compared to other restoration scenarios, these restoration priority areas also displayed the largest combined pressures from dams (75%), mining (67%), and oil and gas (57%) in a 100 km radius. Therefore, forestry emerges as the dominant pressure affecting restoration priority areas focused on ecological integrity and carbon storage, while agriculture and urban development primarily impact priority areas targeting water stability. Nevertheless, unlike conservation scenarios, all restoration priority areas occur across a continuous spectrum of anthropogenic pressures rather than being influenced predominantly by any single one.

Overall, the proximity of future anthropogenic pressures to conservation and restoration priority areas is lower than current pressures (Figure 7). Similar to current patterns of anthropogenic pressures, the conservation priority areas maximizing the protection of threatened species and irrecoverable carbon storage would experience the highest proximity to future natural resource projects, especially mining (i.e., 40% within a 100 km radius). Future mining operations also represented the largest future anthropogenic pressure for all restoration priority areas, especially for 47% of the priority areas aimed at recovering threatened biodiversity with ecological integrity or carbon storage. The potential impacts of mining on these conservation and restoration priority areas are disproportionately concentrated in the high human footprint Mixedwood plains (> 58% within a 100 km radius). Future mining is expected to occur in a 100 km radius of 29% of restoration priority areas recovering threatened biodiversity habitats and surface water stability. However, a slightly larger proportion (31%) of these priority areas are expected to be influenced by future oil and gas operations. Future oil and gas projects pose particularly substantial challenges for biodiversity-water restoration efforts in the Atlantic Maritimes and Prairies. Assuming that current anthropogenic pressures remain in the next decade, these results imply that the mining sector represents the largest growing pressure on both conservation and restoration priority areas.

**Figure 7.**
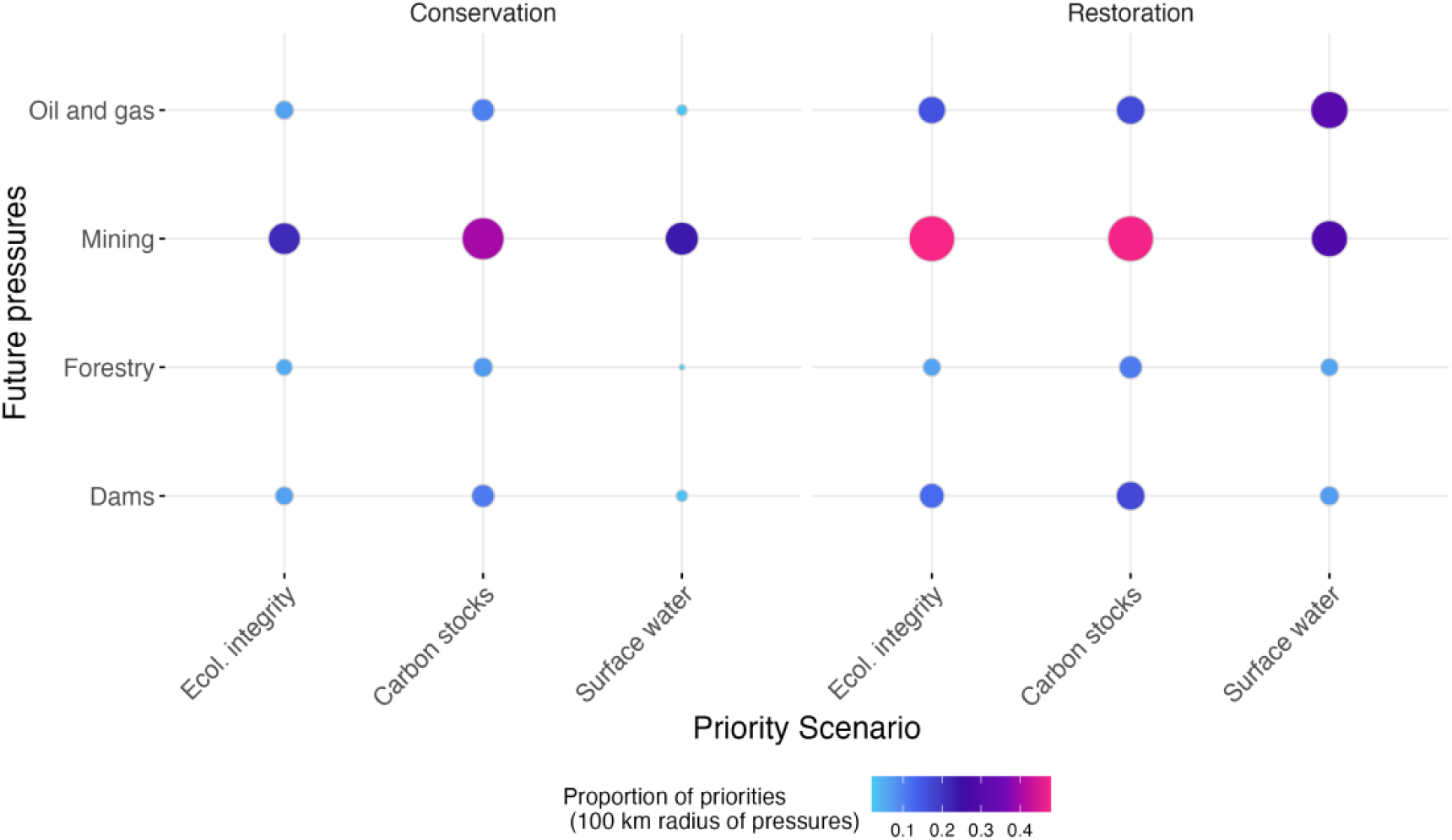
The proportion of conservation and restoration priority areas within a 100 km range of future anthropogenic pressures from major natural resource projects planned in the next ten years.

## Discussion

Our study strengthens a growing body of research aiming to inform the impactful implementation of Nature-based Solutions. Leveraging Reinforcement Learning, we provide an integrated prioritization framework for conservation and restoration actions that maximizes threatened biodiversity protection in synergy with co-benefits from ecological integrity, carbon storage, and water. We test this framework to achieve Canada’s 30×30 conservation and restoration targets stated in the Kunming-Montreal Global Biodiversity Framework. The results exhibit environmental trade-offs along with potential synergies, demonstrating how Artificial Intelligence can help optimize conservation and restoration decisions. Particularly, this integrated prioritization framework reveals combinations of conservation and restoration actions that maximize both biodiversity protection and other environmental co-benefits. The current and future anthropogenic pressures of conservation and restoration scenarios uncover shared socio-economic challenges and coordinated opportunities for implementing effective Nature-based Solutions across government levels and economic sectors.

### Potential pathways to conserve 30% of land

Our results highlight opportunities to reach 30×30 conservation targets that improve biodiversity while maximizing co-benefits. Currently, our results confirm that the existing Protected Areas network in Canada does not effectively protect threatened and non-threatened species (Eckert et al., 2023) and only partially captures the potential co-benefits from carbon storage and water (Mitchell et al., 2021). These results are not exclusive to Canada and reflect a known global trend where Protected Areas are biased to locations with reduced land-use pressures that do not necessarily prioritize biodiversity and other co-benefits outcomes (Joppa & Pfaff, 2009).

To improve Protected Areas outcomes, our results show that prioritizing the protection of threatened biodiversity and irrecoverable carbon stocks yields improvements in biodiversity and multiple co-benefits. Expanding the Protected Areas network with this prioritization approach could increase ecological integrity, carbon storage and surface water stability. Most of these priorities are located in southern provinces, extending across forest-dominated lands in the Boreal ecozones, and the western Montane Cordillera. Without accounting for biodiversity, Mitchell et al. (2021) have already suggested the potential of these ecozones for carbon storage, water provision, and nature-based recreation. Although not assessed in our analysis, these forest lands serve as climate refugia for multiple taxonomic groups, including vertebrates, plants, and invertebrates (Saunders et al., 2023; Stralberg et al., 2020). Our results, therefore, demonstrate that conservation targets regarding biodiversity and ecological integrity can be effectively aligned with avoided land use emissions targets in Nationally Determined Contributions while also achieving water co-benefits.

Our prioritization framework also highlights strategic locations with exceptional conservation value for biodiversity, irrecoverable carbon storage, ecological integrity, and water outcomes. A few conservation priorities in the Atlantic Maritimes, Mixedwood Plains, and Prairies surpass biodiversity and co-benefits outcomes in the Boreal ecozones and Montane Cordillera that could moderately but strategically contribute to conserving 30% of land. These priorities may guarantee the occurrence of climate-resilient species (Eckert et al. 2023) and contain regions under moderate to high anthropogenic pressures with moderate to low conservation actions (Kraus and Hebb 2020).

Other strategic priority locations are ecozones transitions. For example, the peatland dominated James Bay Lowlands (Hudson Plains) and the forested Abitibi Plains (Boreal Shield) in Quebec and Ontario. Similarly, conservation priorities maximizing multiple co-benefits across western Canada would connect the Pacific Maritimes with the Montane Cordillera, Taiga, and Boreal Plains in British Columbia and Alberta. The latter ecozones transition may strategically contribute to the Yellowstone to Yukon corridor (Chester, 2015). Moreover, both ecozone transitions have been identified as either climate corridors or refugia (Saunders et al., 2023; Stralberg et al., 2020). By targeting areas with limited human footprint that maintain the habitat and connectivity of threatened species, these conservation priorities are expected to enhance species flow (Brennan et al., 2022), Protected Areas’ connectivity (Custode et al., 2023), environmental outcomes, and climate resilience.

Although prioritizing threatened species protection and irrecoverable carbon storage may strengthen existing Protected Areas outcomes, such criteria would increase the overlap with existing human pressures. This increase is mostly explained by the current anthropogenic pressure from forestry activities and dams, which has been especially documented in the Boreal ecozones (Wells et al., 2020). The proximity of these pressures to conservation priorities suggests a combination of Protected Areas and Other Effective Area-based Conservation Measures (OECMs) to conserve 30% of land by 2030. OECMs may promote practices in forestry and hydropower generation moderate negative impacts on biodiversity and related co-benefits. For example, privileging retention forestry over conventional practices (Fedrowitz et al., 2014). Given the customary land governance of Indigenous Peoples across low human footprint ecosystems (Artelle et al., 2019; Fa et al., 2020), Indigenous-led OECMs also have the potential to synergize environmental and economic outcomes with cultural revitalization and self-determination (Nitah, 2021; Reed et al., 2021; Vogel et al., 2022).

### Potential pathways to restore 30% of land

The restoration priority scenarios suggest diverse and context-dependent pathways to restore 30% of degraded lands. These restoration scenarios focus on non-intact areas that have lost habitat quality and connectivity for threatened species, along with ecological integrity, carbon stocks, and surface water stability. Contrasting previous frameworks solely focusing on converted lands for cattle and agriculture (Strassburg et al., 2020), our framework’s scenarios suggest complementary restoration pathways in disturbed natural lands and converted lands. The restoration pathway in disturbed natural lands emerges from scenarios where restoring lost biodiversity was prioritized, along with targeted improvements in ecological integrity or irrecoverable carbon stocks. The resulting restoration priority areas are dispersed across forest-dominated Boreal ecozones and the Montane Cordillera, often positioned near conservation priorities.

Beyond confirming previous findings (Currie et al., 2023), our results show that these priority areas have a moderate current human footprint, while still retaining irrecoverable carbon storage and surface water stability. These features make them suitable candidates for restoration actions with short-term benefits. For instance, combining actions within and around remaining forests, such as improved forest management and reforestation (Rayden et al., 2023), could improve ecological integrity and other environmental outcomes. Additionally, restoring and mitigating further carbon stocks losses in these forest lands would result in longer-term climate benefits than restoration in agricultural lands (Matthews et al., 2023).

Similar to conservation priority areas, forestry is the dominant anthropogenic pressure affecting potential restoration priority areas that target recovering threatened biodiversity along with ecological integrity and carbon storage. The Forestry industry has reconfigured forests’ composition and structure through varied harvesting impacts, reducing habitat and connectivity for numerous taxonomic groups (Venier et al., 2014). However, the losses of ecological integrity and carbon storage cannot be uniquely attributed to the direct effects of forestry. North America has experienced the second-largest wildfire-induced losses in timber-producing forests (5.8-6.6 Mha) over the past two decades (Bousfield et al., 2023). Unless the global 2**°**C temperature target above pre-industrial level is achieved, Canada’s projected wildfires are expected to increase by the end of the century (Curasi et al., 2024). Addressing potential climate pressures demand adaptive actions such as the reduction of land use pressures (Rannow et al., 2014; Welch, 2005), fire prescription (Pereira et al., 2012; Wang et al., 2022), translocating or slowing dispersal of species, along with other forms of ecosystem engineering (Lemieux & Scott, 2011; Rannow et al., 2014). Implementing these adaptive actions across restoration priorities in disturbed forested lands will contribute to achieving the 30% restoration target and increasing climate resilience. Furthermore, the proximity of restoration priority areas, conservation priority areas, and forestry represents an opportunity for governments at different levels to coordinate actions with private stakeholders that may close the biodiversity and climate finance gap (Cook-Patton et al., 2021).

The restoration pathway in converted lands resulted from prioritizing areas that have lost habitat quality and connectivity for threatened species, along with surface water stability. Expectedly, these restoration priority areas have moderate to high human footprints in areas dominated by agriculture and urban development, including the Prairies, Mixedwood Plains and Atlantic Maritimes. Previous studies have highlighted biodiversity potential within such ecozones for both conservation (Eckert et al., 2023) and restoration (Currie et al., 2023). More aligned with the latter, our results indicate that these heavily disturbed landscapes are suitable for diverse actions aiming to improve biodiversity and multiple water co-benefits (e.g., water provision, quality, flood reduction). For instance, avoiding further grassland conversion for cropland, maintaining shelterbelts, and increasing tree intercropping and silvopasture are essential actions to reduce agriculture’s anthropogenic pressures (Drever et al., 2021; Keesstra et al., 2018). Combining actions such as green roofs, vegetative swales, and bioretention has been proven effective in urban settings (Huang et al., 2020). Additional actions include riparian tree planting across grasslands, croplands (Scherr & McNeely, 2008), and urban areas (Hutchins et al., 2024). Evidently, small to medium-scale actions alone may not achieve restoration targets. Nevertheless, moderately improving biodiversity and water co-benefits by combining restoration actions can lead to direct benefits for agricultural production (Rey Benayas & Bullock, 2012) and public health in urban areas (van den Bosch & Ode Sang, 2017).

In addition to small and medium-scale actions, our results show the potential for large-scale actions in agricultural and urban landscapes to achieve restoration targets. For instance, wetland and river restoration represent large-scale actions to enhance biodiversity and water co-benefits that simultaneously create new livelihood and recreational opportunities (Souliotis & Voulvoulis, 2022). The significant losses of carbon stocks in the Prairies and Mixedwood plains due to agriculture represent one of the largest opportunities to simultaneously restore carbon and biodiversity in Canada (Currie et al., 2023). However, these large-scale actions carry some caveats. The prairies are not extensive climate refugia or corridors (Eckert et al., 2023; Saunders et al., 2023; Stralberg et al., 2020), and the Mixewood Plains are expected to experience increased climate instability (Herrando-Moraira et al., 2022). Shifting land use may also prove economically unfeasible (Drever et al., 2021; Griscom et al., 2017), constraining the scale of restoration. Addressing climate pressures, while safeguarding food production and human habitation, will require further prioritization scenarios at higher resolutions that account for opportunity costs (Cook-Patton et al., 2021). Consequently, the potential of large-scale restoration in urban and agricultural lands toward the 30% target is highly constrained by social and economic trade-offs. These results suggest that effectively restoring 30% of land will require a combination of targeted actions in natural and transformed ecosystems.

### Future implications of conservation and restoration targets

Current conservation and restoration priority areas face dominant pressures from forestry, agriculture, and urban development, yet different sectors will generate the largest pressure increases in the coming decade. Oil sands extraction in northeastern Alberta and mining along the Ontario-Québec border will directly impact forested priority areas, with immediate effects on threatened biodiversity, ecological integrity, and carbon storage. Mining and energy development will similarly affect restoration priority areas for biodiversity and water stability across the developed Prairies and Mixedwood Plains. Mining impacts extend beyond habitat removal to landscape-scale disturbances through light and chemical pollution, altering biodiversity through complex pathways (Roberts et al., 2022). Moreover, mining will exacerbate existing pressures in areas that have already lost ecological integrity (Sonter et al., 2018). Specific extraction methods may also produce distinct consequences. Water-intensive potash mining expansion in the Prairies threatens water availability for human populations and agriculture (Clark et al., 2017), while pipeline expansion across western ecozones will fragment habitat and alter species composition (Richardson et al., 2017). These results imply that major natural resource projects will expand the baseline of degraded land, undermining the achievement of effective conservation and restoration targets by 2030.

### Study limitations and future directions

Despite developing an integrated framework to achieve conservation and restoration targets, our study has some limitations. First, the complexity of conservation and restoration planning underscores the need for further research at higher resolutions and smaller spatial scales. For instance, high-resolution scenarios could display low-scale conservation priorities in agricultural and urban landscapes that are ruled out in our framework. Second, our framework does not directly account for the demand or immediate benefits from biodiversity and related co-benefits. This limitation is illustrated by surface water stability, which displays areas experiencing flooding or drought (Pekel et al., 2016), but could be complemented by assessing water provision and demand (Mitchell et al., 2021). Similarly, carbon storage co-benefits from NbS in natural or transformed lands would not necessarily result in interchangeable climate benefits. To address disparate climate benefits, Matthews et al. (2023) propose tracking the amount of carbon storage achieved over time and estimating the near-term temperature benefits from that storage. Lastly, while we compare our results with previous work on climate scenarios and assess potential land use conflicts, our prioritization framework does not directly incorporate these assessments. Future prioritization frameworks would benefit from integrating climate and land use scenarios to envision NbS in different shared socio-economic pathways.

### Conclusions

Our study advances Nature-based Solutions implementation through an AI-driven framework that simultaneously addresses the interconnected challenges of biodiversity loss, climate change, and water insecurity. After assessing species habitat quality and connectivity, our framework explores diverse conservation and restoration scenarios to reach Canada’s 30×30 biodiversity targets in synergy with ecological integrity, climate and water co-benefits. Our results indicate that prioritizing the protection of threatened species and irrecoverable carbon storage in forest lands represents an optimal pathway to improve existing Protected Areas outcomes and conserve 30% of land. Effectively restoring 30% of degraded land to protect threatened biodiversity will require a combination of scenarios: restoring ecological integrity and carbon stocks in disturbed forested lands and mitigating the effects of reduced surface water instability in transformed lands. The assessment of anthropogenic pressures reveals that forestry in natural lands and agriculture and urban development in transformed lands would greatly benefit from supporting both restoration and conservation actions. These findings offer complementary insights for environmental decision-making. On the one hand, exploring diverse priority scenarios may reveal strategic conservation and restoration locations that maximize environmental outcomes. On the other hand, assessing current and future anthropogenic pressures suggests targeted and coordinated conservation and restoration actions, policies, and investments that can address environmental and socio-economic trade-offs.

To accelerate the strategic implementation and investments of optimal NbS, our results point to four key governance recommendations that emphasize multi-dimensional coordination. First, NbS for conservation and restoration actions must be interconnected to guarantee long-term outcomes (Wiens & Hobbs, 2015). Second, conservation and restoration actions should be coordinated across different government levels, asserting Indigenous land governance. Part of this administrative coordination will imply balancing economic activities against conservation and restoration priorities. The third dimension of coordination involves the public and private sectors. Traditionally, the former relies on policy interventions for conservation, and the latter invests in restoration (Cook-Patton et al., 2021). Nevertheless, reshaping these traditional roles may close the biodiversity and climate finance gap. Finally, decision-making in the digital age requires coordinating scientific, collective, and artificial intelligence (Luers, 2021) through ethical and human-centered frameworks (Shneiderman, 2021). These transdisciplinary frameworks will require co-developing with regional stakeholders and local rightsholders on which targets, species, co-benefits, actions, scale, data, and model training are most relevant in their specific contexts.

## Acknowledgements

This work has been supported by a Natural Sciences and Engineering Research Council of Canada (NSERC) Alliance grant with Microsoft (ALLPR 571271-21).

C.A. was supported by the Horizon Postdoctoral fellowship at Concordia University.

## Authors Contributions

Conceptualization, C.A., A.L., H.D.M.; Methodology, C.A., H.D.M., M.I.A.P.; Formal Analysis, Investigation, Data Curation, Writing – Original Draft, Visualization, Project Administration C.A.; Writing – Review & Editing, C.A., H.D.M., A.L., A.V.; M.I.A.P.; Supervision, H.D.M.; Funding Acquisition H.D.M., A.L.

