## Supplementary Material for "Maximizing Nature-based Solutions using Artificial Intelligence to align global biodiversity, climate, and water targets"

**Table S1. Datasets used to determine and analyze Conservation and Restoration priority scenarios.**

| Category | Data | Source | Year | Resolution |
|---|---|---|---|---|
| Biodiversity and ecological integrity | Threatened vertebrates (excluding birds) and plant species distribution | (IUCN, 2023) | 2023 | Polygon |
| | Threatened bird species distribution | *(BirdLife International & Handbook of the Birds of the World, 2022)* | | |
| | ESA- CCI Land Cover | (Defourny et al., 2023) | 1992, 2020 | ~ 300 m |
| | Elevation | (Natural Resources Canada, 2011) | 2011 | ~90 m |
| Carbon Stocks | Aboveground and belowground biomass | (Spawn et al., 2020) | 2010 | ~ 300 m |
| | Soil Organic Carbon | (Sothe et al., 2022) | 2015 | ~ 250 m |
| | Irrecoverable Carbon Stocks | (Noon et al., 2022) | 2010 | ~300 m |
| | ESA- CCI Land Cover | (Defourny et al., 2023) | 2010, 2020 | ~ 300 m |
| | Petlands distribution | (Xu et al., 2018) | 2018 | Polygon |
| Surface Water | Surface Water Occurrence Change Intensity | (Pekel et al., 2016) | 1984, 2015 | ~ 30 m |
| Protected Areas | Canadian Protected and Conserved Areas Database | (Environment and Climate Change Canada, 2023) | 2023 | Polygon |
| Ecozones | Terrestrial Ecozones of Canada | (Agriculture and Agri-Food Canada, 2023) | 2023 | Polygon |
| Current Anthropogenic Pressures | Canada's human footprint | (Hirsh-Pearson et al., 2022) | 2016 | ~ 300 m |
| | Built environments (urban, roads, railways) | (Agriculture and Agri-Food Canada, 2020) | 2020 | ~ 30 m |
| | Agriculture and grasslands | (Agriculture and Agri-Food Canada, 2020) | 2020 | ~ 30 m |
| | Oil and gas fields | (Natural Resources Canada, 2020) | 2020 | Point |
| | Mining | (Natural Resources Canada, 2020) | 2020 | Point |
| | Forestry | (Hermosilla et al., 2016) | 2020 | ~ 30 m |
| | Dams and reservoirs | (Lehner et al., 2024) | 2020 | Polygon |

| | | | | |
|---|---|---|---|---|
| Future Anthropogenic Pressures | Major Projects Inventory (Energy, Mining, Forestry) | (Natural Resources Canada, 2024) | 2023 | Point, line |

**Table S2. Biomass and soil organic carbon change estimates from land cover transitions.** *Vegetation land cover classes include forest, wetland, shrubland, grassland, and sparse vegetation. ** Vegetation land cover classes were classified according to their carbon stocks levels as: (forest ~ wetland) > shrubland > sparse vegetation > grassland. *** Mean biomass carbon stocks (ABC) were calculated for all land cover classes in each Canadian Terrestrial Ecozone. ****ABC recovery after 5 years. *****SOC recovery after 40 years.

| Initial land cover | Final land cover | Biomass (ABC) | Soil Organic Carbon (SOC) non-peatlands (0.3 m depth) | Soil Organic Carbon (SOC) peatlands (1m depth) |
|---|---|---|---|---|
| Vegetation*/Agriculture | Bare Land/Urban/Water | Vegetation/Agriculture ABC*0% (Noon et al., 2022) | SOC*0.05*year (Noon et al., 2022) | SOC*0% (Noon et al., 2022) |
| High C Vegetation** | Low C Vegetation** | High ABC Vegetation – Ecozone ABC Mean for Low C Biomass Vegetation*** (Hu et al., 2021) | No change | No change |
| Forest/Shrubland/ Sparse vegetation | Agriculture | Forest /Shrubland/Sparse vegetation ABC – Ecozone ABC Mean for Agriculture (Hu et al., 2021) | SOC -32.19*[1-exp(-years/5.15)] (Poeplau et al., 2011) | SOC*0% (Noon et al., 2022) |
| Grassland | Agriculture | Forest Biomass – Ecozone Mean for Agriculture Biomass (Hu et al., 2021) | SOC -36.11[1*exp(years/2.74)] (Poeplau et al., 2011) | SOC*0% (Noon et al., 2022) |
| Wetland | Shrubland/Grassland | Wetland ABC – Ecozone ABC Mean for Shrubland/Grassland *** (Hu et al., 2021) | SOC -(2.8*years)(Penman et al., 2003) | SOC -(2.8*years) (Penman et al., 2003) |
| Forest/Wetland | Wetland/Forest | No change (Amani et al., 2021; Li et al., 2023). | No change (Amani et al., 2021; Li et al., 2023). | No change (Amani et al., 2021; Li et al., 2023). |

| | | | | |
|---|---|---|---|---|
| Low C Vegetation**/Bare Land/Urban/Water | Boreal Forest/Shrubland/ Sparse Vegetation | ABC + 44.06*ln(years) - 87.534 (Noon et al., 2022) | SOC +1.2*years (Poeplau et al., 2011) | SOC+0.21*years (Poeplau et al., 2011) |
| Low C Vegetation**/Bare Land/Urban/Water | Temperate Lowland Forest/Shrubland/ Sparse Vegetation | ABC + 33.285*ln(years) - 47.14 (Noon et al., 2022) | SOC +1.2*years (Poeplau et al., 2011) | SOC+0.28*years (Poeplau et al., 2011) |
| Low C Vegetation**/Bare Land/Urban/Water | Temperate Mountain Forest/Shrubland/ Sparse Vegetation | ABC + 21.973*ln(years) - 23.6 (Loranty et al., 2014; Noon et al., 2022) | SOC +1.2*years (Poeplau et al., 2011) | SOC+0.28*years (Poeplau et al., 2011) |
| Low C Vegetation**/Bare Land/Urban/Water | Tundra High C Vegetation** | [ABC + (Ecozone ABC Mean for Tundra High C Vegetation – ABC)] / [1 + exp(-2*(years - 2.5))]**** (Rocha & Shaver, 2011) | SOC +1.2*years (Poeplau et al., 2011) | [SOC + (Ecozone Mean SOC – SOC] / (1 + exp(-0.25 * (years - 20)))***** (Loranty et al., 2014) |

| Overlapped Scenarios | Ecozone | Priorties | Threatened species (Ind./100km2) | | Species richness (Ind./100km2) | | Ecological integrity (SHI) | | Carbon stocks (tC/ha) | | Surface Water Change (%) | |
|---|---|---|---|---|---|---|---|---|---|---|---|---|
| | | | x̄ | σ | x̄ | σ | x̄ | σ | x̄ | σ | x̄ | σ |
| Integrity-Carbon-Water | Arctic Cordillera | 1 | 5.00 | NA | 22.00 | NA | 1.00 | NA | 63.02 | NA | 27.17 | NA |
| Integrity-Water | Arctic Cordillera | 19 | 4.58 | 1.84 | 33.89 | 10.96 | 0.89 | 0.08 | 23.78 | 6.89 | 9.61 | 1.66 |
| Carbon-Water | Atlantic Maritime | 15 | 11.80 | 4.13 | 168.73 | 27.38 | 0.61 | 0.10 | 616.41 | 89.98 | 6.89 | 1.65 |
| Integrity-Carbon-Water | Atlantic Maritime | 59 | 6.86 | 2.32 | 126.71 | 29.88 | 0.85 | 0.11 | 612.84 | 91.15 | 8.45 | 1.83 |
| Integrity-Carbon | Atlantic Maritime | 568 | 6.51 | 1.37 | 129.24 | 15.73 | 0.79 | 0.08 | 609.30 | 60.94 | 11.66 | 2.90 |
| Carbon-Water | Boreal Cordillera | 40 | 8.23 | 2.69 | 115.33 | 25.85 | 0.65 | 0.13 | 517.92 | 76.94 | 7.58 | 0.31 |
| Integrity-Water | Boreal Cordillera | 4 | 6.50 | 1.29 | 96.00 | 13.83 | 0.78 | 0.04 | 528.48 | 112.37 | 7.99 | 0.79 |
| Integrity-Carbon | Boreal Cordillera | 884 | 5.46 | 0.95 | 87.04 | 15.76 | 0.85 | 0.09 | 570.29 | 98.67 | 13.87 | 2.93 |
| Integrity-Carbon-Water | Boreal Cordillera | 133 | 5.39 | 1.32 | 85.18 | 15.80 | 0.88 | 0.09 | 538.22 | 67.09 | 8.08 | 1.78 |
| Carbon-Water | Boreal Plains | 649 | 12.31 | 3.41 | 143.60 | 27.67 | 0.60 | 0.16 | 483.61 | 82.23 | 5.32 | 1.45 |
| Integrity-Water | Boreal Plains | 41 | 10.93 | 3.37 | 134.51 | 28.25 | 0.77 | 0.09 | 336.12 | 61.49 | 5.84 | 1.03 |
| Integrity-Carbon-Water | Boreal Plains | 375 | 8.60 | 2.80 | 110.97 | 27.49 | 0.83 | 0.10 | 513.58 | 104.04 | 6.50 | 2.62 |
| Integrity-Carbon | Boreal Plains | 748 | 7.70 | 2.29 | 112.36 | 21.58 | 0.79 | 0.08 | 667.88 | 126.27 | 13.02 | 2.21 |
| Carbon-Water | Boreal Shield | 2412 | 11.67 | 3.32 | 144.00 | 33.70 | 0.61 | 0.17 | 472.54 | 89.90 | 5.29 | 1.30 |
| Integrity-Water | Boreal Shield | 194 | 8.29 | 3.30 | 101.34 | 30.10 | 0.76 | 0.08 | 312.67 | 105.51 | 5.33 | 1.14 |
| Integrity-Carbon-Water | Boreal Shield | 993 | 7.90 | 3.08 | 114.89 | 32.66 | 0.82 | 0.11 | 514.11 | 129.28 | 5.34 | 1.96 |
| Integrity-Carbon | Boreal Shield | 1668 | 6.03 | 2.43 | 102.50 | 34.87 | 0.82 | 0.10 | 585.63 | 122.17 | 10.22 | 1.84 |
| Carbon-Water | Hudson Plains | 37 | 11.70 | 2.13 | 132.59 | 21.06 | 0.60 | 0.08 | 377.08 | 78.30 | 7.60 | 1.37 |
| Integrity-Water | Hudson Plains | 10 | 9.60 | 2.63 | 81.70 | 12.37 | 0.78 | 0.06 | 263.63 | 54.75 | 7.19 | 0.93 |
| Integrity-Carbon-Water | Hudson Plains | 13 | 8.31 | 4.21 | 93.69 | 25.03 | 0.86 | 0.12 | 403.45 | 118.65 | 9.48 | 1.91 |
| Integrity-Carbon | Hudson Plains | 18 | 7.78 | 3.51 | 98.83 | 17.91 | 0.77 | 0.08 | 443.15 | 126.87 | 10.24 | 1.61 |
| Integrity-Carbon | Mixedwood Plains | 13 | 15.00 | 2.08 | 147.23 | 34.53 | 0.79 | 0.15 | 198.81 | 79.08 | 2.54 | 1.70 |
| Integrity-Carbon-Water | Mixedwood Plains | 10 | 14.40 | 3.57 | 150.00 | 42.95 | 0.81 | 0.14 | 250.08 | 208.79 | 2.87 | 1.80 |
| Carbon-Water | Mixedwood Plains | 6 | 12.33 | 2.50 | 118.67 | 10.09 | 0.82 | 0.19 | 355.54 | 298.38 | 3.33 | 0.76 |
| Carbon-Water | Montane Cordillera | 186 | 7.06 | 1.98 | 138.91 | 25.02 | 0.62 | 0.07 | 813.83 | 170.38 | 5.85 | 1.20 |
| Integrity-Water | Montane Cordillera | 7 | 6.57 | 2.15 | 142.86 | 28.38 | 0.76 | 0.07 | 606.15 | 207.05 | 7.11 | 2.24 |
| Integrity-Carbon | Montane Cordillera | 932 | 4.94 | 1.38 | 109.49 | 24.03 | 0.80 | 0.08 | 878.77 | 192.12 | 10.77 | 2.29 |
| Integrity-Carbon-Water | Montane Cordillera | 391 | 4.92 | 1.38 | 111.91 | 24.84 | 0.79 | 0.08 | 906.34 | 181.01 | 6.55 | 1.49 |
| Integrity-Carbon-Water | Northern Arctic | 8 | 6.25 | 2.05 | 23.00 | 6.93 | 0.92 | 0.05 | 61.37 | 16.72 | 10.11 | 10.12 |
| Carbon-Water | Northern Arctic | 5 | 5.80 | 2.95 | 20.60 | 9.24 | 0.65 | 0.01 | 59.78 | 19.54 | 1.14 | 2.56 |
| Integrity-Water | Northern Arctic | 739 | 4.93 | 1.86 | 24.30 | 6.03 | 0.88 | 0.12 | 40.78 | 14.66 | 6.78 | 1.81 |
| Integrity-Carbon | Northern Arctic | 21 | 4.33 | 0.48 | 17.90 | 2.02 | 0.85 | 0.13 | 66.72 | 22.41 | 24.47 | 3.28 |
| Carbon-Water | Pacific Maritime | 33 | 6.45 | 3.37 | 90.27 | 27.43 | 0.62 | 0.04 | 1336.69 | 187.82 | 5.88 | 0.71 |
| Integrity-Carbon | Pacific Maritime | 88 | 4.92 | 1.48 | 108.24 | 24.14 | 0.77 | 0.07 | 1188.50 | 209.88 | 13.00 | 3.30 |
| Integrity-Carbon-Water | Pacific Maritime | 38 | 3.47 | 0.92 | 63.53 | 15.63 | 0.77 | 0.08 | 1398.93 | 165.91 | 5.92 | 0.05 |
| Integrity-Water | Prairies | 82 | 10.60 | 2.71 | 133.50 | 23.92 | 0.90 | 0.11 | 115.14 | 98.91 | 10.27 | 2.50 |
| Integrity-Carbon-Water | Prairies | 154 | 9.77 | 1.23 | 119.39 | 16.32 | 0.95 | 0.06 | 88.46 | 67.68 | 12.47 | 2.42 |
| Integrity-Carbon | Prairies | 29 | 9.41 | 2.21 | 137.93 | 27.56 | 0.86 | 0.12 | 315.55 | 285.60 | 13.11 | 0.98 |
| Carbon-Water | Prairies | 17 | 9.35 | 0.79 | 110.53 | 5.20 | 1.02 | 0.01 | 65.94 | 8.77 | 12.32 | 2.56 |
| Carbon-Water | Southern Arctic | 7 | 8.57 | 1.72 | 54.29 | 15.55 | 0.70 | 0.14 | 187.14 | 17.12 | 7.31 | 1.44 |
| Integrity-Carbon-Water | Southern Arctic | 58 | 7.41 | 1.08 | 45.93 | 6.69 | 0.82 | 0.07 | 171.47 | 29.70 | 7.22 | 1.05 |
| Integrity-Water | Southern Arctic | 785 | 6.13 | 1.44 | 40.33 | 6.79 | 0.81 | 0.10 | 109.86 | 49.53 | 6.86 | 1.30 |
| Integrity-Carbon | Southern Arctic | 7 | 5.57 | 1.13 | 31.86 | 12.10 | 0.92 | 0.07 | 93.91 | 61.22 | 14.38 | 5.02 |
| Integrity-Carbon-Water | Taiga Cordillera | 1 | 7.00 | NA | 58.00 | NA | 0.88 | NA | 191.06 | NA | 9.97 | NA |
| Integrity-Water | Taiga Cordillera | 4 | 6.75 | 0.50 | 61.50 | 1.91 | 0.75 | 0.00 | 222.89 | 10.89 | 9.97 | 0.00 |
| Integrity-Carbon | Taiga Cordillera | 40 | 5.55 | 0.81 | 79.98 | 13.21 | 0.81 | 0.09 | 405.76 | 94.86 | 11.65 | 1.17 |
| Carbon-Water | Taiga Plains | 129 | 10.45 | 2.73 | 124.35 | 27.07 | 0.62 | 0.13 | 414.43 | 74.29 | 5.56 | 1.17 |
| Integrity-Carbon-Water | Taiga Plains | 235 | 7.25 | 2.01 | 92.96 | 18.03 | 0.86 | 0.09 | 468.92 | 96.35 | 6.68 | 1.72 |
| Integrity-Carbon | Taiga Plains | 588 | 6.61 | 1.30 | 96.00 | 15.16 | 0.82 | 0.09 | 582.59 | 108.49 | 14.65 | 4.18 |
| Integrity-Water | Taiga Plains | 148 | 5.93 | 2.31 | 70.54 | 19.86 | 0.80 | 0.09 | 280.09 | 63.87 | 5.51 | 1.07 |
| Carbon-Water | Taiga Shield | 153 | 10.78 | 2.67 | 105.63 | 29.89 | 0.61 | 0.04 | 312.77 | 106.45 | 5.82 | 0.58 |
| Integrity-Carbon-Water | Taiga Shield | 92 | 8.78 | 3.06 | 90.02 | 26.12 | 0.75 | 0.07 | 289.67 | 86.01 | 6.43 | 1.62 |
| Integrity-Water | Taiga Shield | 157 | 7.11 | 2.60 | 70.47 | 26.33 | 0.75 | 0.07 | 204.46 | 98.79 | 6.68 | 1.71 |
| Integrity-Carbon | Taiga Shield | 207 | 3.75 | 1.53 | 56.71 | 11.11 | 0.82 | 0.12 | 402.79 | 137.30 | 11.70 | 2.25 |

**Table S3.** Environmental outcomes of conservation scenarios overlapped priorities in Canadian terrestrial ecozones. x̄ and σ symbolize the average and standard deviation, respectively.

| Overlapped Scenarios | Ecozone | Priorties | Threatened species | | Species richness (Ind./100km2) | | Ecological integrity (SHI) | | Carbon stocks (tC/ha) | | Surface Water Change (%) | |
|---|---|---|---|---|---|---|---|---|---|---|---|---|
| | | | x̄ | σ | x̄ | σ | x̄ | σ | x̄ | σ | x̄ | σ |
| Integrity-Carbon | Atlantic Maritime | 110 | 10.75 | 4.28 | 155.39 | 30.57 | 0.55 | 0.05 | 505.98 | 69.91 | 9.75 | 1.71 |
| Integrity-Carbon-Water | Atlantic Maritime | 11 | 8.09 | 3.33 | 138.55 | 19.68 | 0.62 | 0.10 | 554.85 | 54.96 | 11.78 | 3.18 |
| Integrity-Water | Atlantic Maritime | 26 | 11.73 | 4.16 | 158.96 | 24.58 | 0.56 | 0.05 | 495.50 | 121.48 | 11.65 | 2.29 |
| Carbon-Water | Boreal Cordillera | 40 | 6.23 | 1.40 | 93.43 | 18.90 | 0.73 | 0.12 | 514.65 | 107.77 | 13.88 | 2.60 |
| Integrity-Carbon | Boreal Cordillera | 73 | 7.71 | 2.00 | 112.88 | 20.41 | 0.50 | 0.04 | 436.08 | 108.31 | 11.68 | 3.73 |
| Integrity-Carbon-Water | Boreal Cordillera | 4 | 8.25 | 1.71 | 124.00 | 25.88 | 0.55 | 0.01 | 544.43 | 137.18 | 14.26 | 3.86 |
| Integrity-Water | Boreal Cordillera | 5 | 8.80 | 2.17 | 119.80 | 20.24 | 0.54 | 0.02 | 533.92 | 86.47 | 14.26 | 2.31 |
| Carbon-Water | Boreal Plains | 384 | 8.46 | 2.98 | 126.68 | 27.28 | 0.73 | 0.15 | 618.68 | 178.34 | 13.54 | 1.72 |
| Integrity-Carbon | Boreal Plains | 924 | 11.11 | 3.31 | 145.77 | 26.01 | 0.50 | 0.05 | 555.35 | 132.46 | 9.98 | 3.73 |
| Integrity-Carbon-Water | Boreal Plains | 58 | 12.36 | 4.24 | 159.17 | 37.21 | 0.55 | 0.02 | 569.62 | 139.21 | 13.67 | 1.70 |
| Integrity-Water | Boreal Plains | 45 | 12.18 | 3.21 | 161.33 | 31.60 | 0.56 | 0.02 | 500.03 | 219.50 | 14.50 | 2.77 |
| Carbon-Water | Boreal Shield | 109 | 10.28 | 5.07 | 148.13 | 50.12 | 0.71 | 0.15 | 632.25 | 134.06 | 11.89 | 1.79 |
| Integrity-Carbon | Boreal Shield | 1444 | 10.53 | 3.52 | 131.86 | 43.25 | 0.51 | 0.04 | 463.47 | 107.10 | 8.60 | 2.85 |
| Integrity-Carbon-Water | Boreal Shield | 25 | 16.00 | 4.62 | 191.80 | 42.01 | 0.56 | 0.10 | 592.16 | 143.85 | 11.88 | 1.97 |
| Integrity-Water | Boreal Shield | 32 | 13.59 | 4.45 | 166.63 | 48.75 | 0.55 | 0.02 | 544.56 | 187.00 | 11.49 | 1.87 |
| Carbon-Water | Hudson Plains | 3 | 14.00 | 2.65 | 103.33 | 13.80 | 0.65 | 0.09 | 316.66 | 42.76 | 12.79 | 2.90 |
| Integrity-Carbon | Hudson Plains | 28 | 10.89 | 3.54 | 100.89 | 20.67 | 0.49 | 0.05 | 252.32 | 44.21 | 9.33 | 1.86 |
| Integrity-Carbon-Water | Hudson Plains | 6 | 15.50 | 2.35 | 118.50 | 18.04 | 0.56 | 0.01 | 306.93 | 89.91 | 14.21 | 2.62 |
| Integrity-Water | Hudson Plains | 3 | 17.00 | 1.00 | 133.00 | 22.61 | 0.56 | 0.01 | 147.91 | NA | 11.48 | 0.36 |
| Carbon-Water | Mixedwood Plains | 57 | 15.63 | 3.17 | 198.77 | 25.36 | 0.67 | 0.11 | 525.55 | 133.20 | 12.04 | 1.58 |
| Integrity-Carbon | Mixedwood Plains | 44 | 16.50 | 4.69 | 202.66 | 38.90 | 0.56 | 0.06 | 486.85 | 156.92 | 6.52 | 4.23 |
| Integrity-Carbon-Water | Mixedwood Plains | 12 | 21.50 | 2.58 | 246.17 | 17.64 | 0.57 | 0.02 | 505.85 | 136.51 | 12.33 | 1.48 |
| Integrity-Water | Mixedwood Plains | 36 | 19.81 | 4.14 | 229.67 | 35.64 | 0.56 | 0.03 | 423.87 | 118.46 | 12.32 | 1.53 |
| Carbon-Water | Montane Cordillera | 130 | 5.81 | 1.52 | 116.21 | 21.23 | 0.72 | 0.09 | 752.30 | 162.10 | 12.29 | 1.72 |
| Integrity-Carbon | Montane Cordillera | 308 | 7.10 | 1.77 | 152.10 | 27.42 | 0.53 | 0.04 | 788.57 | 158.82 | 8.41 | 2.50 |
| Integrity-Carbon-Water | Montane Cordillera | 10 | 8.30 | 1.64 | 133.50 | 13.25 | 0.55 | 0.03 | 705.90 | 67.12 | 13.25 | 2.49 |
| Integrity-Water | Montane Cordillera | 5 | 9.00 | 1.00 | 141.20 | 10.55 | 0.54 | 0.03 | 735.30 | 148.67 | 13.78 | 2.11 |
| Carbon-Water | Pacific Maritime | 10 | 5.70 | 2.21 | 110.50 | 19.73 | 0.73 | 0.05 | 1042.50 | 105.01 | 13.09 | 2.60 |
| Integrity-Carbon | Pacific Maritime | 102 | 7.57 | 2.80 | 120.54 | 28.31 | 0.53 | 0.03 | 1149.18 | 217.41 | 8.88 | 3.00 |
| Integrity-Carbon-Water | Pacific Maritime | 1 | 10.00 | NA | 185.00 | NA | 0.56 | NA | 1047.54 | NA | 10.47 | NA |
| Integrity-Water | Pacific Maritime | 3 | 11.00 | 1.73 | 149.33 | 15.01 | 0.55 | 0.01 | 916.33 | 214.54 | 13.63 | 3.60 |
| Carbon-Water | Prairies | 47 | 11.06 | 2.27 | 145.91 | 28.23 | 0.81 | 0.12 | 209.00 | 195.84 | 14.08 | 2.06 |
| Integrity-Carbon | Prairies | 9 | 14.22 | 4.41 | 184.33 | 31.06 | 0.64 | 0.13 | 237.11 | 149.56 | 10.54 | 3.29 |
| Integrity-Carbon-Water | Prairies | 14 | 13.14 | 2.82 | 174.21 | 22.23 | 0.70 | 0.08 | 196.73 | 133.85 | 13.43 | 1.24 |
| Integrity-Water | Prairies | 10 | 13.30 | 2.98 | 167.70 | 24.99 | 0.59 | 0.09 | 157.12 | 46.85 | 14.36 | 3.19 |
| Integrity-Carbon | Taiga Cordillera | 8 | 6.50 | 0.53 | 91.88 | 8.82 | 0.50 | 0.02 | 196.39 | 51.67 | 11.68 | 2.38 |
| Carbon-Water | Taiga Plains | 26 | 6.73 | 1.28 | 102.15 | 13.20 | 0.78 | 0.09 | 626.45 | 89.72 | 16.32 | 2.48 |
| Integrity-Carbon | Taiga Plains | 72 | 9.56 | 2.73 | 119.32 | 24.85 | 0.50 | 0.04 | 496.20 | 145.90 | 11.37 | 3.73 |
| Integrity-Carbon-Water | Taiga Plains | 11 | 9.00 | 2.79 | 124.73 | 26.43 | 0.54 | 0.02 | 574.29 | 102.71 | 15.21 | 2.76 |
| Integrity-Water | Taiga Plains | 12 | 9.67 | 3.14 | 130.25 | 25.52 | 0.55 | 0.01 | 514.40 | 168.69 | 14.74 | 2.52 |
| Carbon-Water | Taiga Shield | 1 | 3.00 | NA | 63.00 | NA | 0.75 | NA | 287.19 | NA | 14.18 | NA |
| Integrity-Carbon | Taiga Shield | 46 | 7.93 | 2.33 | 82.07 | 15.95 | 0.52 | 0.04 | 219.86 | 50.84 | 7.91 | 1.40 |
| Integrity-Water | Taiga Shield | 1 | 6.00 | NA | 85.00 | NA | 0.54 | NA | 164.57 | NA | 11.07 | NA |

**Table S4.** Environmental outcomes of restoration scenarios overlapped priorities in Canadian terrestrial ecozones. x̄ and σ symbolize the average and standard deviation, respectively.

| Conservation scenario | Anthrop. pressure | Ecozone | Distance (km) |  | Priorities (n) | Priorities in a 100 km range of pressure (%) |
|---|---|---|---|---|---|---|
|  |  |  | x̄ | σ |  |  |
| Ecological Integrity | Agriculture | Atlantic Maritime | 885.46 | 701.39 | 1649.00 | 0.17 |
| Ecological Integrity | Agriculture | Boreal Plains | 1520.11 | 562.88 | 312.00 | 0.01 |
| Ecological Integrity | Agriculture | Boreal Shield | 753.81 | 726.37 | 8734.00 | 0.26 |
| Ecological Integrity | Agriculture | Hudson Plains | 1708.17 | 465.91 | 640.00 | NA |
| Ecological Integrity | Agriculture | Mixedwood Plains | 104.19 | 130.80 | 1799.00 | 0.70 |
| Ecological Integrity | Agriculture | Montane Cordillera | 1968.51 | 380.32 | 36.00 | NA |
| Ecological Integrity | Agriculture | Prairies | 1387.97 | 646.71 | 2349.00 | 0.03 |
| Ecological Integrity | Dams | Atlantic Maritime | 549.71 | 503.31 | 1649.00 | 0.20 |
| Ecological Integrity | Dams | Boreal Plains | 985.81 | 505.82 | 312.00 | 0.03 |
| Ecological Integrity | Dams | Boreal Shield | 477.44 | 513.92 | 8734.00 | 0.28 |
| Ecological Integrity | Dams | Hudson Plains | 1147.69 | 508.68 | 640.00 | 0.02 |
| Ecological Integrity | Dams | Mixedwood Plains | 68.24 | 59.63 | 1799.00 | 0.81 |
| Ecological Integrity | Dams | Montane Cordillera | 1346.21 | 481.20 | 36.00 | NA |
| Ecological Integrity | Dams | Prairies | 906.28 | 541.69 | 2349.00 | 0.04 |
| Ecological Integrity | Forestry | Atlantic Maritime | 318.74 | 423.74 | 1649.00 | 0.57 |
| Ecological Integrity | Forestry | Boreal Plains | 706.32 | 463.34 | 312.00 | 0.17 |
| Ecological Integrity | Forestry | Boreal Shield | 277.61 | 427.30 | 8734.00 | 0.64 |
| Ecological Integrity | Forestry | Hudson Plains | 866.02 | 500.10 | 640.00 | 0.14 |
| Ecological Integrity | Forestry | Mixedwood Plains | 13.55 | 48.18 | 1799.00 | 0.97 |
| Ecological Integrity | Forestry | Montane Cordillera | 1086.65 | 576.71 | 36.00 | 0.17 |
| Ecological Integrity | Forestry | Prairies | 608.96 | 511.85 | 2349.00 | 0.28 |
| Ecological Integrity | Mining | Atlantic Maritime | 259.53 | 189.14 | 1649.00 | 0.21 |
| Ecological Integrity | Mining | Boreal Plains | 435.69 | 215.20 | 312.00 | 0.09 |
| Ecological Integrity | Mining | Boreal Shield | 239.28 | 183.02 | 8734.00 | 0.26 |
| Ecological Integrity | Mining | Hudson Plains | 448.27 | 214.41 | 640.00 | 0.07 |
| Ecological Integrity | Mining | Mixedwood Plains | 95.71 | 60.89 | 1799.00 | 0.61 |
| Ecological Integrity | Mining | Montane Cordillera | 709.00 | 193.43 | 36.00 | NA |
| Ecological Integrity | Mining | Prairies | 376.97 | 229.50 | 2349.00 | 0.10 |
| Ecological Integrity | Oil and gas | Atlantic Maritime | 845.89 | 700.65 | 1649.00 | 0.16 |
| Ecological Integrity | Oil and gas | Boreal Plains | 1164.11 | 700.04 | 312.00 | 0.05 |
| Ecological Integrity | Oil and gas | Boreal Shield | 785.46 | 670.24 | 8734.00 | 0.16 |
| Ecological Integrity | Oil and gas | Hudson Plains | 1262.34 | 641.58 | 640.00 | 0.02 |
| Ecological Integrity | Oil and gas | Mixedwood Plains | 481.27 | 242.10 | 1799.00 | 0.08 |
| Ecological Integrity | Oil and gas | Montane Cordillera | 1164.53 | 572.31 | 36.00 | NA |
| Ecological Integrity | Oil and gas | Prairies | 1128.09 | 724.93 | 2349.00 | 0.08 |
| Ecological Integrity | Urban | Atlantic Maritime | 898.64 | 684.73 | 1649.00 | 0.14 |
| Ecological Integrity | Urban | Boreal Plains | 1527.29 | 551.00 | 312.00 | 0.01 |
| Ecological Integrity | Urban | Boreal Shield | 762.07 | 714.02 | 8734.00 | 0.23 |
| Ecological Integrity | Urban | Hudson Plains | 1718.14 | 453.42 | 640.00 | NA |
| Ecological Integrity | Urban | Mixedwood Plains | 90.89 | 116.55 | 1799.00 | 0.77 |
| Ecological Integrity | Urban | Montane Cordillera | 1982.93 | 385.00 | 36.00 | NA |
| Ecological Integrity | Urban | Prairies | 1395.74 | 625.46 | 2349.00 | 0.01 |

**Table S5. The distance of conservation priorities maximizing threatened biodiversity protection and ecological integrity to current anthropogenic pressures across ecozones.**

| Conservation scenario | Anthrop. pressure | Ecozone | Distance (km) | | Priorities (n) | Priorities in a 100 km range of pressure (%) |
|---|---|---|---|---|---|---|
| | | | x̄ | σ | | |
| Irrecoverable Carbon Stocks | Agriculture | Atlantic Maritime | 220.38 | 202.47 | 1625.00 | 0.38 |
| Irrecoverable Carbon Stocks | Agriculture | Boreal Plains | 378.60 | 337.06 | 312.00 | 0.26 |
| Irrecoverable Carbon Stocks | Agriculture | Boreal Shield | 213.05 | 230.30 | 8673.00 | 0.41 |
| Irrecoverable Carbon Stocks | Agriculture | Hudson Plains | 555.74 | 639.09 | 640.00 | 0.18 |
| Irrecoverable Carbon Stocks | Agriculture | Mixedwood Plains | 84.86 | 74.01 | 1799.00 | 0.72 |
| Irrecoverable Carbon Stocks | Agriculture | Montane Cordillera | 774.19 | 201.80 | 24.00 | NA |
| Irrecoverable Carbon Stocks | Agriculture | Prairies | 317.49 | 266.93 | 2267.00 | 0.27 |
| Irrecoverable Carbon Stocks | Agriculture | NA | 187.77 | 199.09 | 1052.00 | 0.49 |
| Irrecoverable Carbon Stocks | Dams | Atlantic Maritime | 131.91 | 86.79 | 1625.00 | 0.43 |
| Irrecoverable Carbon Stocks | Dams | Boreal Plains | 175.85 | 115.52 | 312.00 | 0.29 |
| Irrecoverable Carbon Stocks | Dams | Boreal Shield | 126.09 | 117.76 | 8673.00 | 0.50 |
| Irrecoverable Carbon Stocks | Dams | Hudson Plains | 356.56 | 482.55 | 640.00 | 0.14 |
| Irrecoverable Carbon Stocks | Dams | Mixedwood Plains | 52.48 | 36.09 | 1799.00 | 0.91 |
| Irrecoverable Carbon Stocks | Dams | Montane Cordillera | 182.68 | 107.14 | 24.00 | 0.25 |
| Irrecoverable Carbon Stocks | Dams | Prairies | 166.61 | 111.81 | 2267.00 | 0.34 |
| Irrecoverable Carbon Stocks | Forestry | Atlantic Maritime | 12.02 | 21.97 | 1625.00 | 0.99 |
| Irrecoverable Carbon Stocks | Forestry | Boreal Plains | 17.90 | 48.52 | 312.00 | 0.98 |
| Irrecoverable Carbon Stocks | Forestry | Boreal Shield | 16.35 | 72.53 | 8673.00 | 0.98 |
| Irrecoverable Carbon Stocks | Forestry | Hudson Plains | 163.05 | 470.42 | 640.00 | 0.90 |
| Irrecoverable Carbon Stocks | Forestry | Mixedwood Plains | 9.23 | 34.70 | 1799.00 | 0.98 |
| Irrecoverable Carbon Stocks | Forestry | Montane Cordillera | 32.33 | 55.46 | 24.00 | 0.96 |
| Irrecoverable Carbon Stocks | Forestry | Prairies | 16.07 | 38.16 | 2267.00 | 0.99 |
| Irrecoverable Carbon Stocks | Mining | Atlantic Maritime | 154.14 | 95.82 | 1625.00 | 0.34 |
| Irrecoverable Carbon Stocks | Mining | Boreal Plains | 200.69 | 121.64 | 312.00 | 0.24 |
| Irrecoverable Carbon Stocks | Mining | Boreal Shield | 144.12 | 101.40 | 8673.00 | 0.40 |
| Irrecoverable Carbon Stocks | Mining | Hudson Plains | 260.54 | 222.16 | 640.00 | 0.16 |
| Irrecoverable Carbon Stocks | Mining | Mixedwood Plains | 83.42 | 47.45 | 1799.00 | 0.67 |
| Irrecoverable Carbon Stocks | Mining | Montane Cordillera | 162.15 | 147.10 | 24.00 | 0.50 |
| Irrecoverable Carbon Stocks | Mining | Prairies | 176.16 | 113.47 | 2267.00 | 0.31 |
| Irrecoverable Carbon Stocks | Oil and gas | Atlantic Maritime | 422.46 | 306.21 | 1625.00 | 0.20 |
| Irrecoverable Carbon Stocks | Oil and gas | Boreal Plains | 326.57 | 267.21 | 312.00 | 0.26 |
| Irrecoverable Carbon Stocks | Oil and gas | Boreal Shield | 446.94 | 286.24 | 8673.00 | 0.17 |
| Irrecoverable Carbon Stocks | Oil and gas | Hudson Plains | 361.99 | 471.80 | 640.00 | 0.37 |
| Irrecoverable Carbon Stocks | Oil and gas | Mixedwood Plains | 492.18 | 167.91 | 1799.00 | 0.03 |
| Irrecoverable Carbon Stocks | Oil and gas | Montane Cordillera | 398.71 | 100.93 | 24.00 | NA |
| Irrecoverable Carbon Stocks | Oil and gas | Prairies | 342.03 | 279.54 | 2267.00 | 0.26 |
| Irrecoverable Carbon Stocks | Urban | Atlantic Maritime | 222.42 | 208.82 | 1625.00 | 0.36 |
| Irrecoverable Carbon Stocks | Urban | Boreal Plains | 428.29 | 355.90 | 312.00 | 0.15 |
| Irrecoverable Carbon Stocks | Urban | Boreal Shield | 213.80 | 240.24 | 8673.00 | 0.43 |
| Irrecoverable Carbon Stocks | Urban | Hudson Plains | 608.31 | 635.04 | 640.00 | 0.10 |
| Irrecoverable Carbon Stocks | Urban | Mixedwood Plains | 73.84 | 51.01 | 1799.00 | 0.77 |
| Irrecoverable Carbon Stocks | Urban | Montane Cordillera | 855.87 | 200.28 | 24.00 | NA |
| Irrecoverable Carbon Stocks | Urban | Prairies | 347.73 | 282.31 | 2267.00 | 0.21 |

**Table S6. The distance of conservation priorities maximizing threatened biodiversity protection and irrecoverable carbon storage to current anthropogenic pressures across ecozones.**

| Conservation scenario | Anthrop. pressure | Ecozone | Distance (km) x̄ | Distance (km) σ | Priorities (n) | Priorities in a 100 km range of pressure (%) |
|---|---|---|---|---|---|---|
| Water Surface Stability | Agriculture | Atlantic Maritime | 660.07 | 453.94 | 1625.00 | 0.10 |
| Water Surface Stability | Agriculture | Boreal Plains | 958.26 | 426.78 | 286.00 | 0.03 |
| Water Surface Stability | Agriculture | Boreal Shield | 568.13 | 448.73 | 8593.00 | 0.13 |
| Water Surface Stability | Agriculture | Hudson Plains | 1125.08 | 341.02 | 582.00 | NA |
| Water Surface Stability | Agriculture | Mixedwood Plains | 127.55 | 120.66 | 1799.00 | 0.55 |
| Water Surface Stability | Agriculture | Montane Cordillera | 1651.91 | 561.97 | 24.00 | NA |
| Water Surface Stability | Agriculture | Prairies | 982.50 | 501.79 | 2260.00 | 0.02 |
| Water Surface Stability | Dams | Atlantic Maritime | 343.32 | 293.54 | 1625.00 | 0.18 |
| Water Surface Stability | Dams | Boreal Plains | 495.06 | 330.97 | 286.00 | 0.03 |
| Water Surface Stability | Dams | Boreal Shield | 295.15 | 281.50 | 8593.00 | 0.25 |
| Water Surface Stability | Dams | Hudson Plains | 608.82 | 311.83 | 582.00 | 0.03 |
| Water Surface Stability | Dams | Mixedwood Plains | 73.80 | 58.07 | 1799.00 | 0.78 |
| Water Surface Stability | Dams | Montane Cordillera | 1106.39 | 549.65 | 24.00 | NA |
| Water Surface Stability | Dams | Prairies | 540.07 | 413.15 | 2260.00 | 0.06 |
| Water Surface Stability | Forestry | Atlantic Maritime | 132.82 | 216.37 | 1625.00 | 0.67 |
| Water Surface Stability | Forestry | Boreal Plains | 266.16 | 282.78 | 286.00 | 0.39 |
| Water Surface Stability | Forestry | Boreal Shield | 111.63 | 199.57 | 8593.00 | 0.74 |
| Water Surface Stability | Forestry | Hudson Plains | 341.74 | 274.69 | 582.00 | 0.23 |
| Water Surface Stability | Forestry | Mixedwood Plains | 12.70 | 42.89 | 1799.00 | 0.97 |
| Water Surface Stability | Forestry | Montane Cordillera | 849.01 | 638.05 | 24.00 | 0.04 |
| Water Surface Stability | Forestry | Prairies | 298.23 | 383.03 | 2260.00 | 0.42 |
| Water Surface Stability | Mining | Atlantic Maritime | 216.76 | 143.68 | 1625.00 | 0.20 |
| Water Surface Stability | Mining | Boreal Plains | 294.26 | 184.80 | 286.00 | 0.14 |
| Water Surface Stability | Mining | Boreal Shield | 206.60 | 135.47 | 8593.00 | 0.21 |
| Water Surface Stability | Mining | Hudson Plains | 320.16 | 194.41 | 582.00 | 0.11 |
| Water Surface Stability | Mining | Mixedwood Plains | 98.06 | 64.31 | 1799.00 | 0.58 |
| Water Surface Stability | Mining | Montane Cordillera | 631.24 | 280.01 | 24.00 | NA |
| Water Surface Stability | Mining | Prairies | 288.77 | 203.21 | 2260.00 | 0.13 |
| Water Surface Stability | Oil and gas | Atlantic Maritime | 651.36 | 410.79 | 1625.00 | 0.07 |
| Water Surface Stability | Oil and gas | Boreal Plains | 694.01 | 494.74 | 286.00 | 0.07 |
| Water Surface Stability | Oil and gas | Boreal Shield | 634.35 | 405.74 | 8593.00 | 0.07 |
| Water Surface Stability | Oil and gas | Hudson Plains | 774.98 | 500.37 | 582.00 | 0.04 |
| Water Surface Stability | Oil and gas | Mixedwood Plains | 562.55 | 213.39 | 1799.00 | 0.03 |
| Water Surface Stability | Oil and gas | Montane Cordillera | 986.19 | 548.48 | 24.00 | NA |
| Water Surface Stability | Oil and gas | Prairies | 791.89 | 490.41 | 2260.00 | 0.04 |
| Water Surface Stability | Urban | Atlantic Maritime | 654.14 | 445.33 | 1625.00 | 0.10 |
| Water Surface Stability | Urban | Boreal Plains | 968.96 | 418.94 | 286.00 | 0.02 |
| Water Surface Stability | Urban | Boreal Shield | 555.85 | 443.06 | 8593.00 | 0.14 |
| Water Surface Stability | Urban | Hudson Plains | 1138.53 | 323.78 | 582.00 | NA |
| Water Surface Stability | Urban | Mixedwood Plains | 102.38 | 98.54 | 1799.00 | 0.67 |
| Water Surface Stability | Urban | Montane Cordillera | 1666.62 | 566.32 | 24.00 | NA |
| Water Surface Stability | Urban | Prairies | 981.77 | 495.17 | 2260.00 | 0.02 |

**Table S7. The distance of conservation priorities maximizing threatened biodiversity protection and water surface stability to current anthropogenic pressures across ecozones.**

| Conservation scenario | Anthrop. pressure | Ecozone | Distance (km) | | Priorities (n) | Priorities in a 100 km range of pressure (%) |
|---|---|---|---|---|---|---|
| | | | x̄ | σ | | |
| Ecological Integrity | Agriculture | Atlantic Maritime | 194.90 | 325.83 | 469.00 | 0.63 |
| Ecological Integrity | Agriculture | Boreal Plains | 502.82 | 231.01 | 8.00 | 0.13 |
| Ecological Integrity | Agriculture | Boreal Shield | 150.75 | 232.71 | 3772.00 | 0.60 |
| Ecological Integrity | Agriculture | Mixedwood Plains | 78.21 | 99.40 | 1784.00 | 0.69 |
| Ecological Integrity | Agriculture | Prairies | 562.11 | 74.03 | 8.00 | NA |
| Ecological Integrity | Dams | Atlantic Maritime | 136.12 | 126.68 | 469.00 | 0.48 |
| Ecological Integrity | Dams | Boreal Plains | 182.95 | 122.89 | 8.00 | 0.25 |
| Ecological Integrity | Dams | Boreal Shield | 111.10 | 91.47 | 3772.00 | 0.53 |
| Ecological Integrity | Dams | Mixedwood Plains | 62.70 | 46.71 | 1784.00 | 0.85 |
| Ecological Integrity | Dams | Prairies | 197.42 | 148.39 | 8.00 | 0.38 |
| Ecological Integrity | Forestry | Atlantic Maritime | 9.95 | 34.23 | 469.00 | 0.99 |
| Ecological Integrity | Forestry | Boreal Plains | 27.62 | 33.24 | 8.00 | 0.88 |
| Ecological Integrity | Forestry | Boreal Shield | 8.63 | 41.77 | 3772.00 | 0.99 |
| Ecological Integrity | Forestry | Mixedwood Plains | 7.90 | 22.72 | 1784.00 | 0.99 |
| Ecological Integrity | Forestry | Prairies | 36.76 | 30.57 | 8.00 | 0.88 |
| Ecological Integrity | Mining | Atlantic Maritime | 151.70 | 105.55 | 469.00 | 0.38 |
| Ecological Integrity | Mining | Boreal Plains | 171.48 | 160.18 | 8.00 | 0.50 |
| Ecological Integrity | Mining | Boreal Shield | 133.10 | 94.26 | 3772.00 | 0.43 |
| Ecological Integrity | Mining | Mixedwood Plains | 74.65 | 52.61 | 1784.00 | 0.72 |
| Ecological Integrity | Mining | Prairies | 219.41 | 156.19 | 8.00 | 0.38 |
| Ecological Integrity | Oil and gas | Atlantic Maritime | 306.05 | 329.63 | 469.00 | 0.39 |
| Ecological Integrity | Oil and gas | Boreal Plains | 391.81 | 195.80 | 8.00 | 0.13 |
| Ecological Integrity | Oil and gas | Boreal Shield | 385.52 | 307.14 | 3772.00 | 0.29 |
| Ecological Integrity | Oil and gas | Mixedwood Plains | 484.20 | 215.08 | 1784.00 | 0.07 |
| Ecological Integrity | Oil and gas | Prairies | 345.59 | 144.96 | 8.00 | NA |
| Ecological Integrity | Urban | Atlantic Maritime | 212.53 | 322.71 | 469.00 | 0.58 |
| Ecological Integrity | Urban | Boreal Plains | 555.10 | 243.20 | 8.00 | 0.13 |
| Ecological Integrity | Urban | Boreal Shield | 159.32 | 244.19 | 3772.00 | 0.61 |
| Ecological Integrity | Urban | Mixedwood Plains | 64.56 | 84.58 | 1784.00 | 0.82 |
| Ecological Integrity | Urban | Prairies | 592.85 | 89.94 | 8.00 | NA |

**Table S8. The distance of restoration priorities maximizing threatened biodiversity protection and ecological integrity to current anthropogenic pressures across ecozones.**

| Conservation scenario | Anthrop. pressure | Ecozone | Distance (km) | | Priorities (n) | Priorities in a 100 km range of pressure (%) |
|---|---|---|---|---|---|---|
| | | | x̄ | σ | | |
| Irrecoverable Carbon Stocks | Agriculture | Atlantic Maritime | 90.56 | 119.60 | 515.00 | 0.73 |
| Irrecoverable Carbon Stocks | Agriculture | Boreal Plains | 140.99 | 189.89 | 18.00 | 0.67 |
| Irrecoverable Carbon Stocks | Agriculture | Boreal Shield | 114.12 | 185.39 | 4144.00 | 0.72 |
| Irrecoverable Carbon Stocks | Agriculture | Mixedwood Plains | 59.32 | 78.65 | 1787.00 | 0.79 |
| Irrecoverable Carbon Stocks | Agriculture | Prairies | 201.92 | 187.05 | 50.00 | 0.50 |
| Irrecoverable Carbon Stocks | Dams | Atlantic Maritime | 108.74 | 75.11 | 515.00 | 0.53 |
| Irrecoverable Carbon Stocks | Dams | Boreal Plains | 124.77 | 57.61 | 18.00 | 0.56 |
| Irrecoverable Carbon Stocks | Dams | Boreal Shield | 96.15 | 69.35 | 4144.00 | 0.60 |
| Irrecoverable Carbon Stocks | Dams | Mixedwood Plains | 57.55 | 41.04 | 1787.00 | 0.87 |
| Irrecoverable Carbon Stocks | Dams | Prairies | 179.23 | 96.16 | 50.00 | 0.26 |
| Irrecoverable Carbon Stocks | Forestry | Atlantic Maritime | 5.25 | 8.59 | 515.00 | 1.00 |
| Irrecoverable Carbon Stocks | Forestry | Boreal Plains | 8.99 | 12.46 | 18.00 | 1.00 |
| Irrecoverable Carbon Stocks | Forestry | Boreal Shield | 5.68 | 15.42 | 4144.00 | 1.00 |
| Irrecoverable Carbon Stocks | Forestry | Mixedwood Plains | 8.17 | 24.50 | 1787.00 | 0.98 |
| Irrecoverable Carbon Stocks | Forestry | Prairies | 13.38 | 18.84 | 50.00 | 0.98 |
| Irrecoverable Carbon Stocks | Mining | Atlantic Maritime | 123.11 | 94.68 | 515.00 | 0.50 |
| Irrecoverable Carbon Stocks | Mining | Boreal Plains | 222.95 | 90.18 | 18.00 | 0.17 |
| Irrecoverable Carbon Stocks | Mining | Boreal Shield | 111.73 | 78.04 | 4144.00 | 0.52 |
| Irrecoverable Carbon Stocks | Mining | Mixedwood Plains | 70.73 | 49.38 | 1787.00 | 0.76 |
| Irrecoverable Carbon Stocks | Mining | Prairies | 243.01 | 133.64 | 50.00 | 0.16 |
| Irrecoverable Carbon Stocks | Oil and gas | Atlantic Maritime | 243.81 | 246.79 | 515.00 | 0.38 |
| Irrecoverable Carbon Stocks | Oil and gas | Boreal Plains | 221.42 | 103.76 | 18.00 | 0.06 |
| Irrecoverable Carbon Stocks | Oil and gas | Boreal Shield | 332.12 | 264.25 | 4144.00 | 0.26 |
| Irrecoverable Carbon Stocks | Oil and gas | Mixedwood Plains | 445.10 | 228.11 | 1787.00 | 0.08 |
| Irrecoverable Carbon Stocks | Oil and gas | Prairies | 152.98 | 144.46 | 50.00 | 0.48 |
| Irrecoverable Carbon Stocks | Urban | Atlantic Maritime | 105.32 | 122.38 | 515.00 | 0.71 |
| Irrecoverable Carbon Stocks | Urban | Boreal Plains | 167.90 | 195.84 | 18.00 | 0.67 |
| Irrecoverable Carbon Stocks | Urban | Boreal Shield | 119.49 | 196.18 | 4144.00 | 0.74 |
| Irrecoverable Carbon Stocks | Urban | Mixedwood Plains | 50.42 | 55.44 | 1787.00 | 0.89 |
| Irrecoverable Carbon Stocks | Urban | Prairies | 259.93 | 180.97 | 50.00 | 0.32 |

**Table S9. The distance of restoration priorities maximizing threatened biodiversity protection and irrecoverable carbon storage to current anthropogenic pressures across ecozones.**

| Conservation scenario | Anthrop. pressure | Ecozone | Distance (km) | | Priorities (n) | Priorities in a 100 km range of pressure (%) |
|---|---|---|---|---|---|---|
| | | | x̄ | σ | | |
| Water Surface Stability | Agriculture | Atlantic Maritime | 87.25 | 267.43 | 540.00 | 0.87 |
| Water Surface Stability | Agriculture | Boreal Plains | 143.43 | 162.59 | 18.00 | 0.56 |
| Water Surface Stability | Agriculture | Boreal Shield | 51.74 | 144.41 | 4144.00 | 0.93 |
| Water Surface Stability | Agriculture | Mixedwood Plains | 25.12 | 47.08 | 1788.00 | 0.98 |
| Water Surface Stability | Agriculture | Prairies | 102.10 | 71.44 | 50.00 | 0.50 |
| Water Surface Stability | Dams | Atlantic Maritime | 94.93 | 100.55 | 540.00 | 0.64 |
| Water Surface Stability | Dams | Boreal Plains | 208.01 | 64.03 | 18.00 | 0.06 |
| Water Surface Stability | Dams | Boreal Shield | 83.60 | 67.10 | 4144.00 | 0.70 |
| Water Surface Stability | Dams | Mixedwood Plains | 47.16 | 33.77 | 1788.00 | 0.93 |
| Water Surface Stability | Dams | Prairies | 184.07 | 74.10 | 50.00 | 0.18 |
| Water Surface Stability | Forestry | Atlantic Maritime | 58.69 | 90.88 | 540.00 | 0.77 |
| Water Surface Stability | Forestry | Boreal Plains | 6.81 | 7.19 | 18.00 | 1.00 |
| Water Surface Stability | Forestry | Boreal Shield | 83.50 | 98.61 | 4144.00 | 0.66 |
| Water Surface Stability | Forestry | Mixedwood Plains | 76.66 | 101.21 | 1788.00 | 0.70 |
| Water Surface Stability | Forestry | Prairies | 4.65 | 3.13 | 50.00 | 1.00 |
| Water Surface Stability | Mining | Atlantic Maritime | 97.79 | 75.56 | 540.00 | 0.64 |
| Water Surface Stability | Mining | Boreal Plains | 225.51 | 115.55 | 18.00 | 0.17 |
| Water Surface Stability | Mining | Boreal Shield | 90.78 | 59.25 | 4144.00 | 0.67 |
| Water Surface Stability | Mining | Mixedwood Plains | 78.58 | 46.94 | 1788.00 | 0.73 |
| Water Surface Stability | Mining | Prairies | 219.84 | 120.02 | 50.00 | 0.18 |
| Water Surface Stability | Oil and gas | Atlantic Maritime | 101.65 | 162.01 | 540.00 | 0.74 |
| Water Surface Stability | Oil and gas | Boreal Plains | 231.96 | 311.02 | 18.00 | 0.61 |
| Water Surface Stability | Oil and gas | Boreal Shield | 118.92 | 155.69 | 4144.00 | 0.64 |
| Water Surface Stability | Oil and gas | Mixedwood Plains | 217.14 | 187.20 | 1788.00 | 0.38 |
| Water Surface Stability | Oil and gas | Prairies | 84.17 | 121.47 | 50.00 | 0.78 |
| Water Surface Stability | Urban | Atlantic Maritime | 117.02 | 276.27 | 540.00 | 0.82 |
| Water Surface Stability | Urban | Boreal Plains | 200.96 | 152.54 | 18.00 | 0.50 |
| Water Surface Stability | Urban | Boreal Shield | 83.38 | 156.90 | 4144.00 | 0.87 |
| Water Surface Stability | Urban | Mixedwood Plains | 42.09 | 48.38 | 1788.00 | 0.97 |
| Water Surface Stability | Urban | Prairies | 151.17 | 109.59 | 50.00 | 0.42 |

**Table S10. The distance of restoration priorities maximizing threatened biodiversity protection and irrecoverable carbon storage to current anthropogenic pressures across ecozones.**

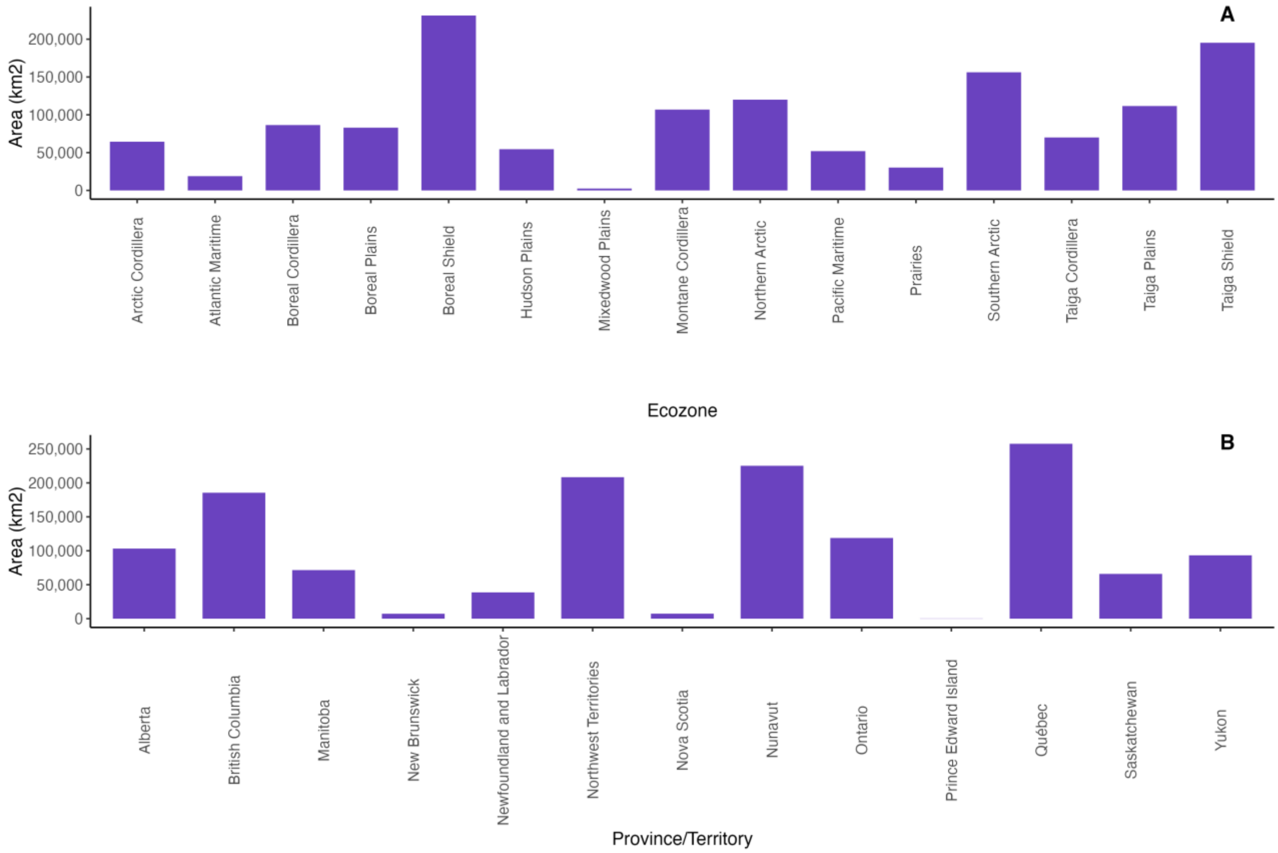


**Figure S1. The land surface extent of Protected Areas in Canada. A. Area distribution across Canadian Terrestrial Ecozones. B. Area distribution across Provinces and Territories.**

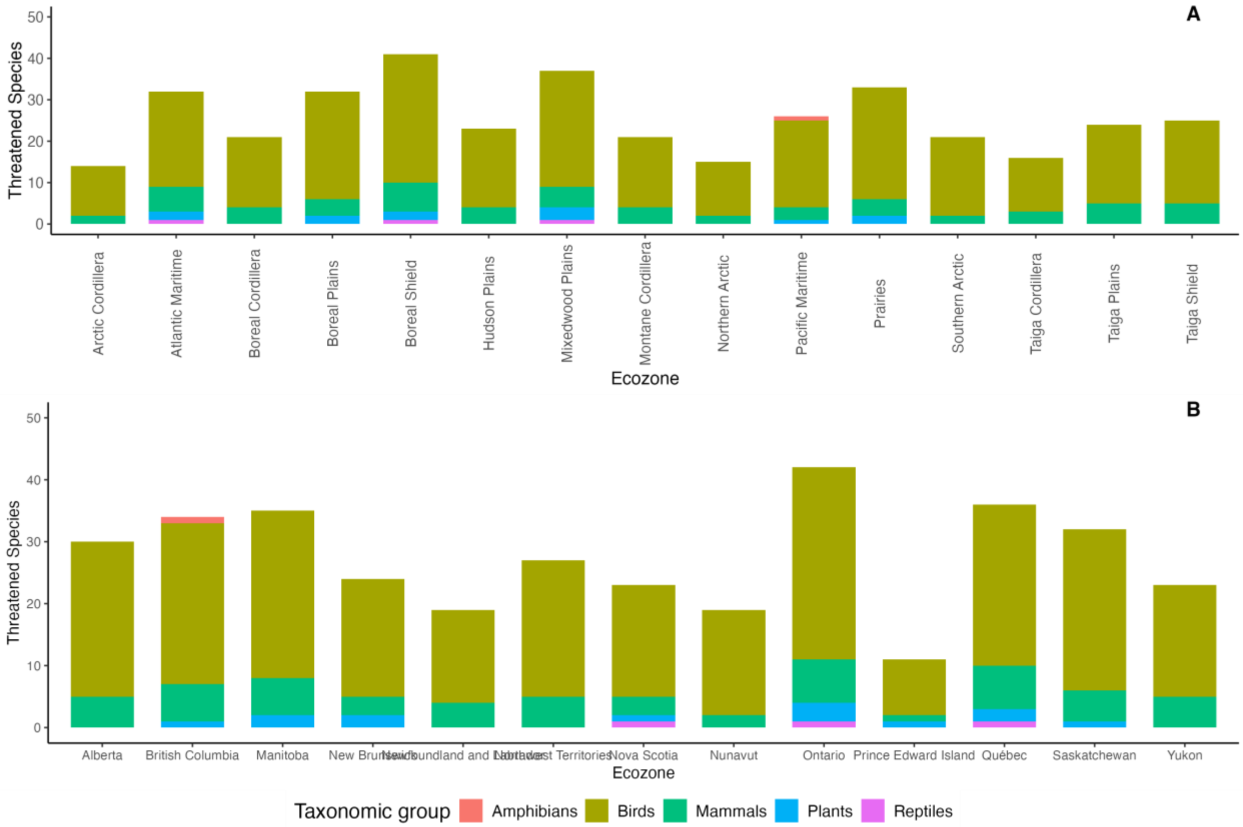


**Figure S2. The distribution of threatened vertebrate and plant species in Canada. A. Distribution across Canadian Terrestrial Ecozones. B. A. Distribution across Provinces and Territories.**

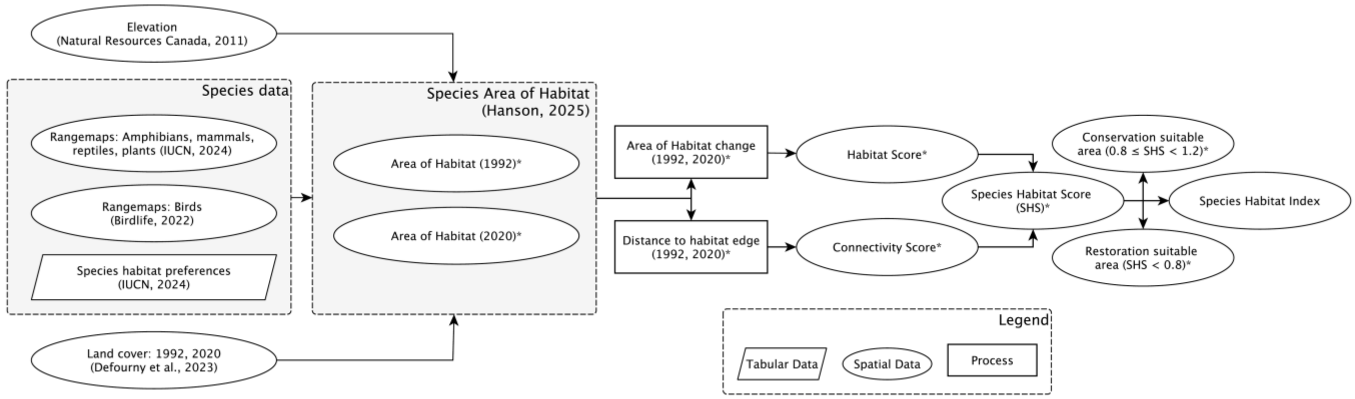


**Figure S3. Workflow explaining the estimation of Species Habitat Scores and the Species Habitat Index. *Processes and information for a single species.**

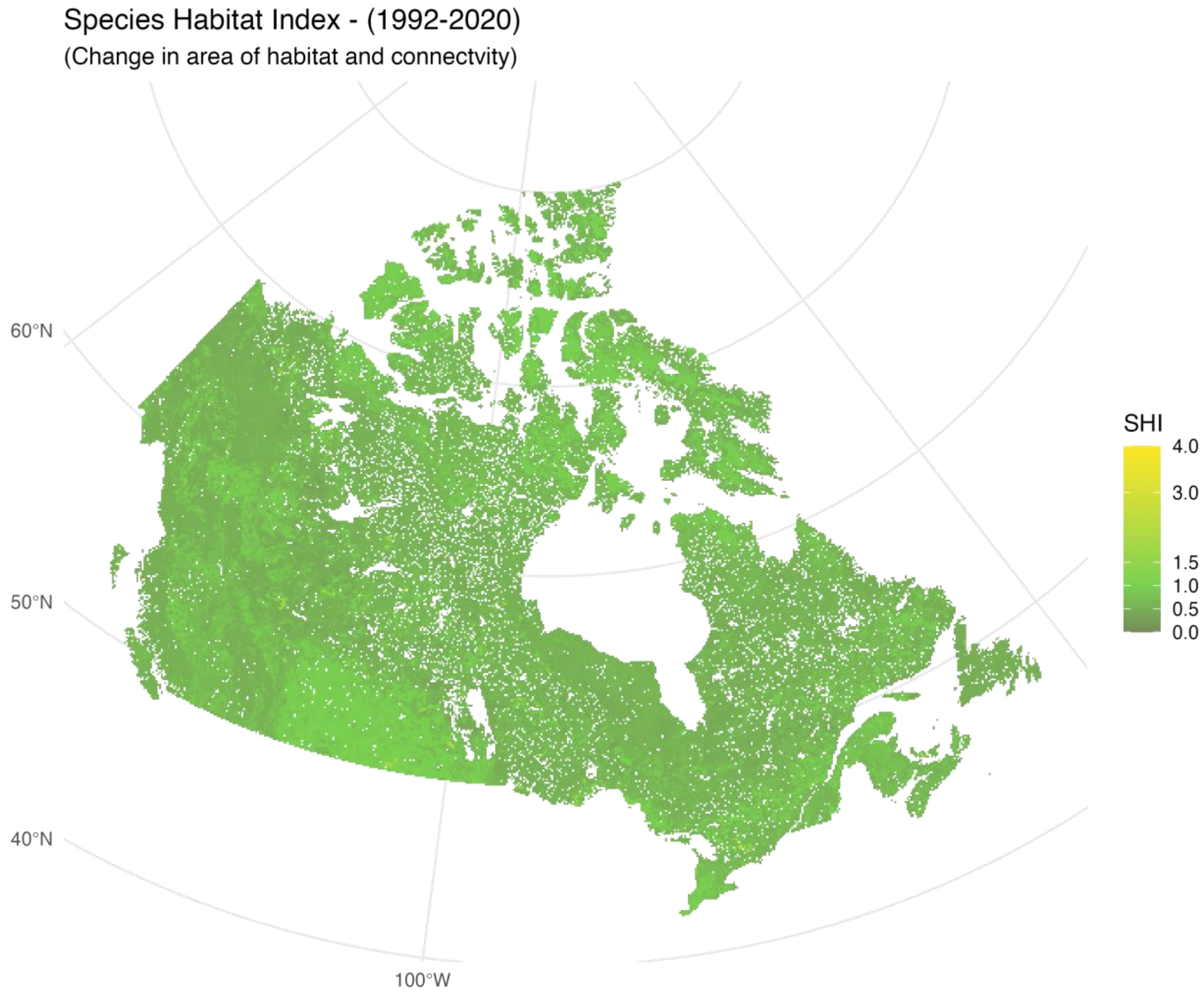


**Figure S4. Species Habitat Index in 10x10km planning units across Canada.**

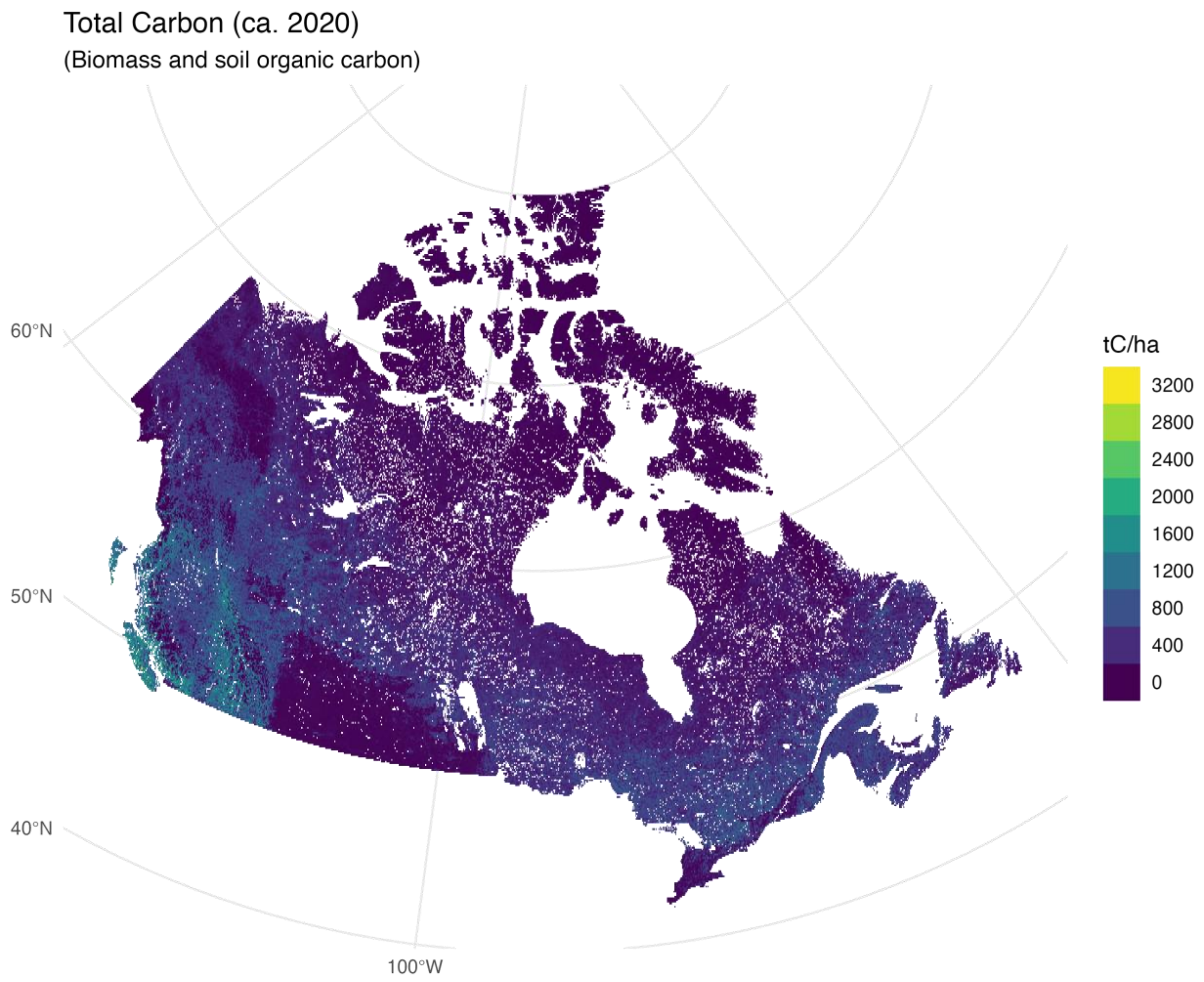


**Figure S5. The spatial distribution of total carbon stocks in Canada (ca. 2020).** Total carbon stocks represent the sum of aboveground and belowground biomass with soil organic carbon.

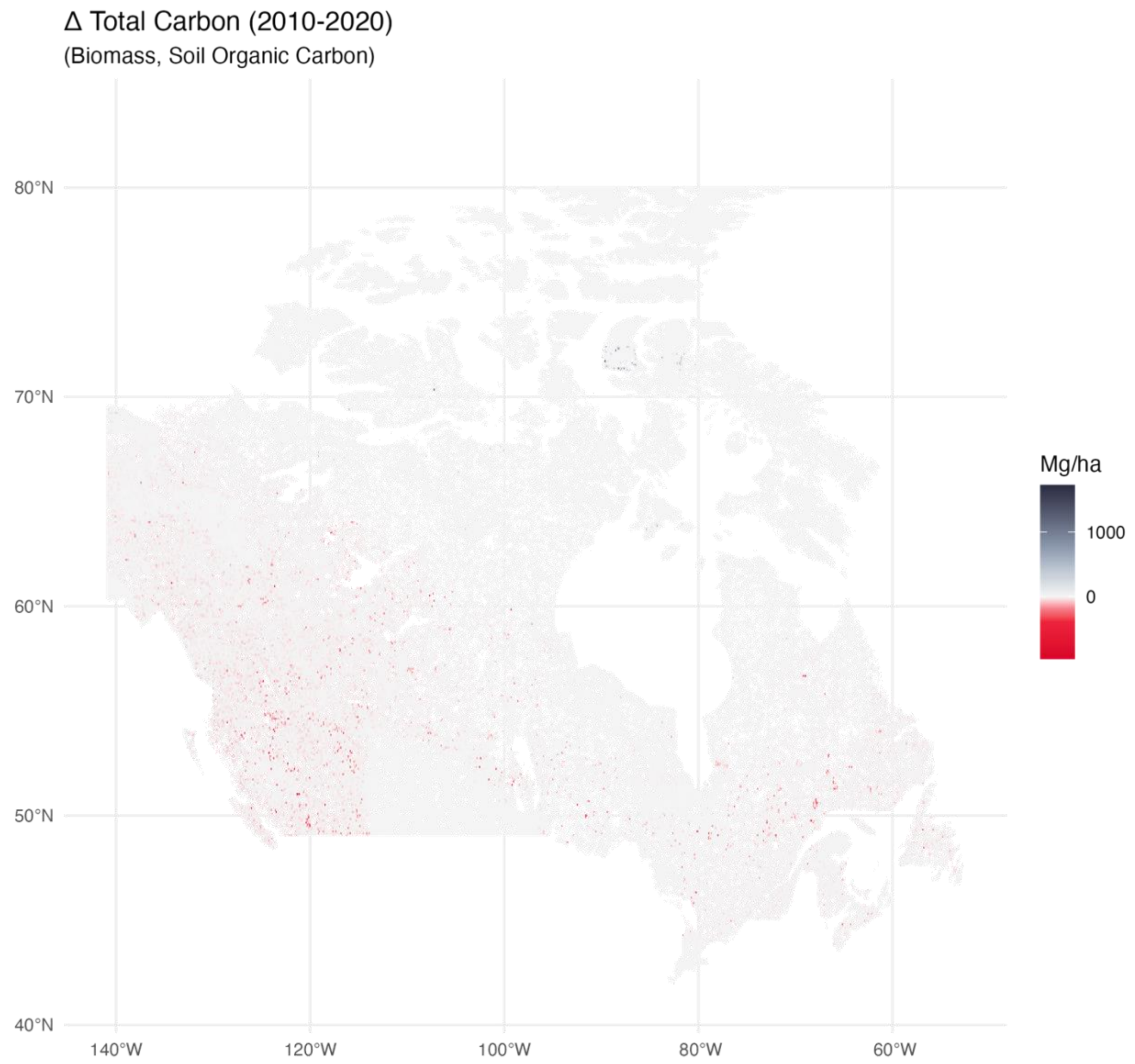


**Figure S6. Total carbon stocks changes (2010-2020).**

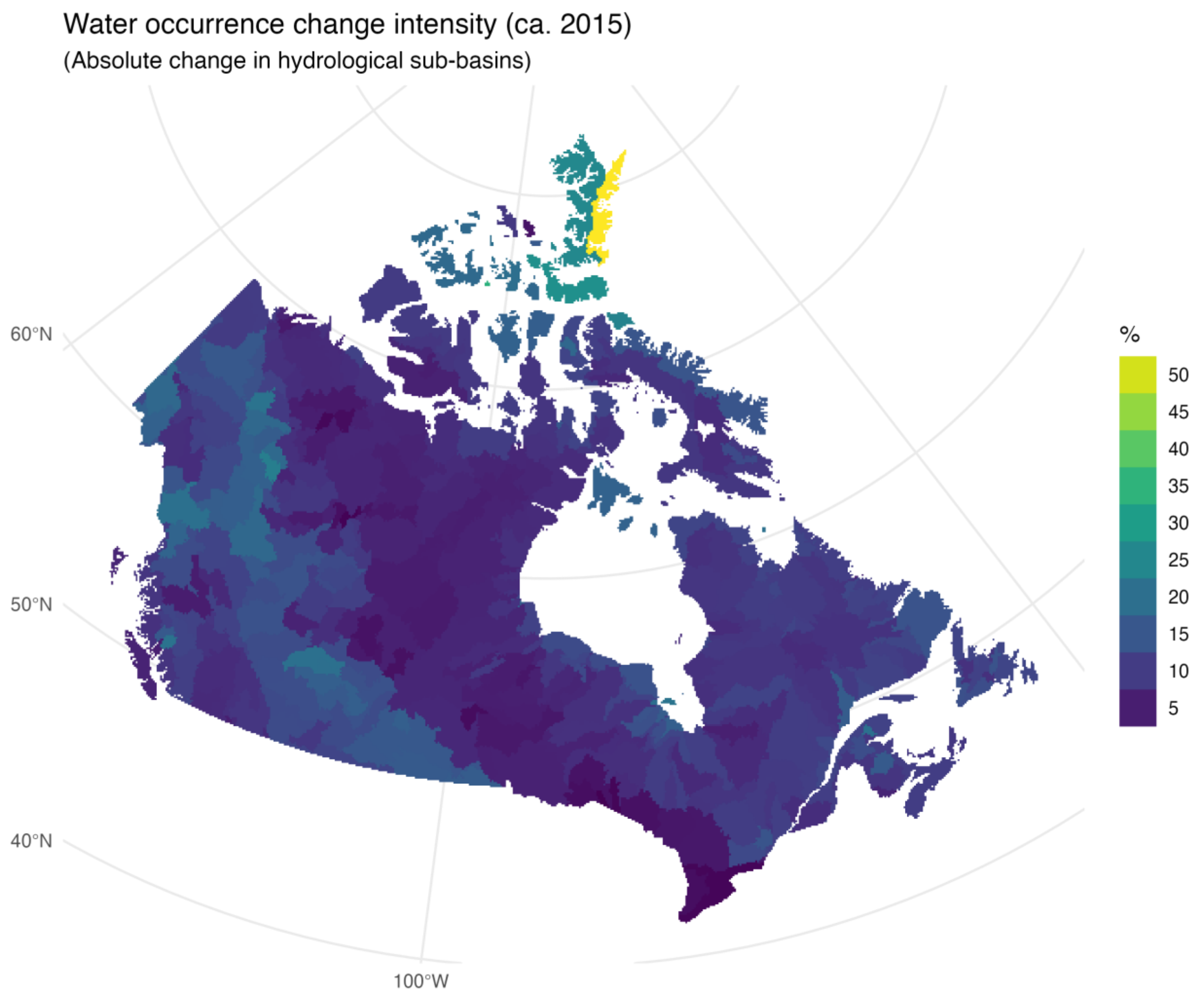


**Figure S7. Average changes in surface water in hydrological sub-basins in Canada (1984-2015).**

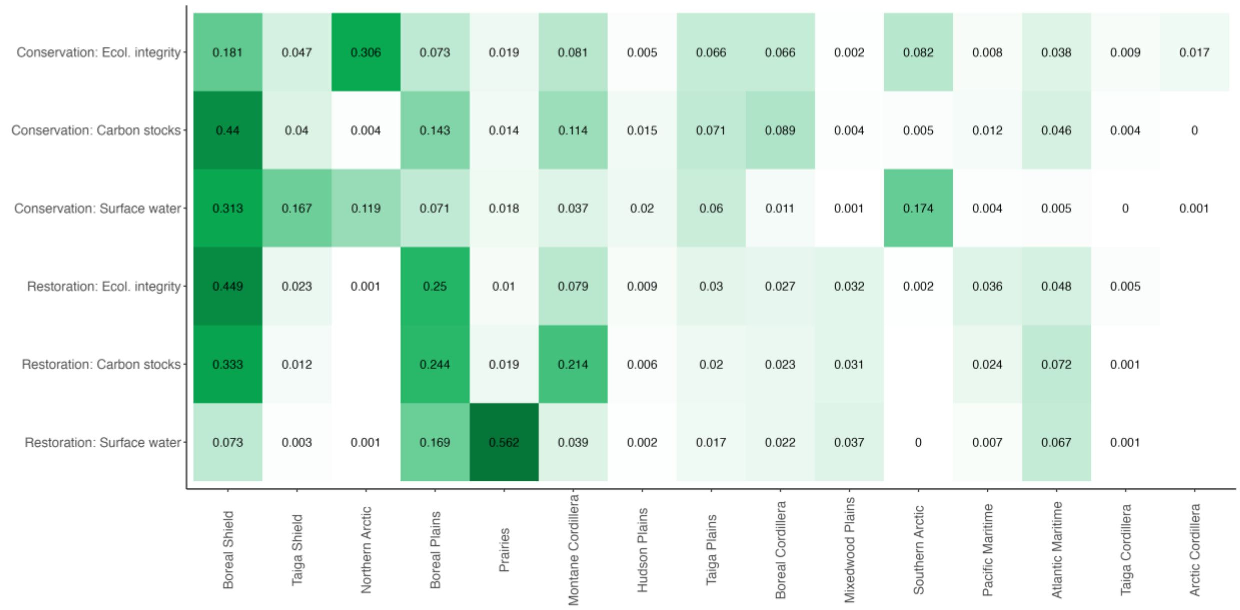


***Figure S8.*** **The distribution of Conservation and Restoration Priority Scenarios across Canadian Terrestrial Ecozones**. All scenarios aim to maximize the occurrence of threatened species. The conservation scenarios additionally prioritize the occurrence of high Ecological Integrity, Irrecoverable Carbon Stocks, and Water Surface Stability. The restoration scenarios additionally prioritize the occurrence of low ecological intactness, irrecoverable carbon stocks losses, and water surface instability. The Scenarios Overlaps represent the co-occurrence of conservation or restoration priorities in at least two scenarios.

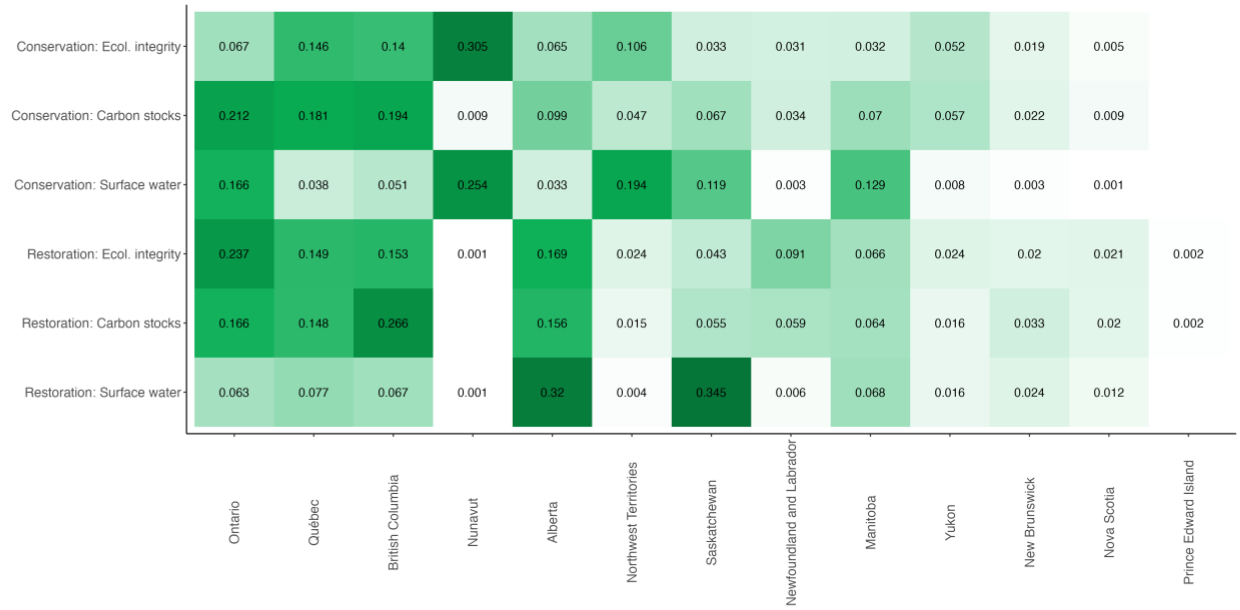

**Figure S9. The distribution of Conservation and Restoration Priority Scenarios across Canadian provinces**. All scenarios aim to maximize the occurrence of threatened species. The conservation scenarios additionally prioritize the occurrence of high Ecological Intactness, Irrecoverable Carbon Stocks, and Water Surface Stability. The restoration scenarios additionally prioritize the occurrence of low ecological intactness, irrecoverable carbon stocks losses, and water surface instability. The Scenarios Overlaps represent the co-occurrence of conservation or restoration priorities in at least two scenarios.

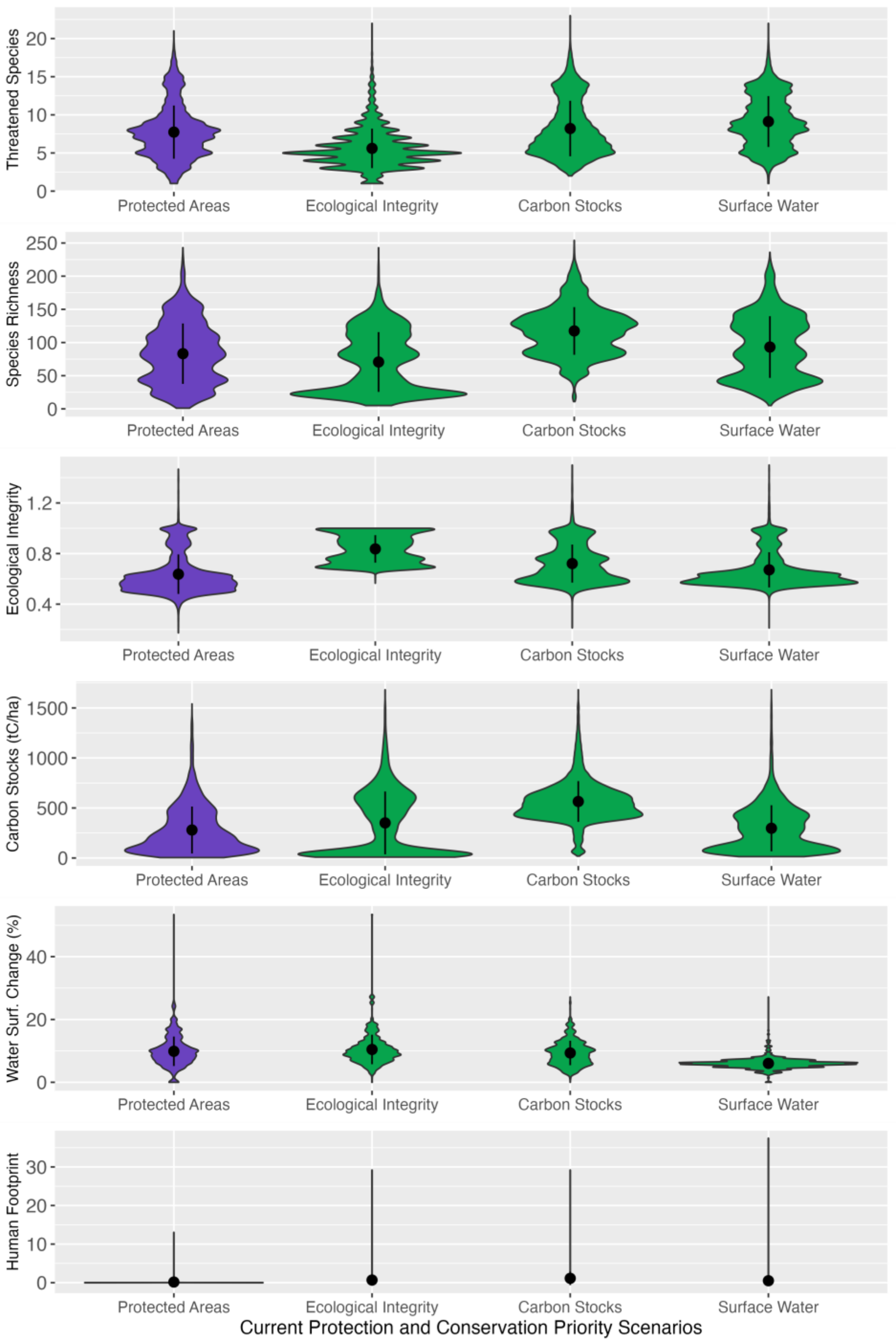


**Figure S10. The distribution of threatened species, species richness, ecological integrity, carbon stocks, absolute surface water change, and human footprint in existing Protected Areas and Conservation Priority Scenarios.** The conservation scenarios aim to maximize the occurrence of threatened species with co-benefits emerging from Ecological Integrity, Irrecoverable Carbon Stocks, and Water Surface Stability. The points and bars refer to the mean and standard deviation.

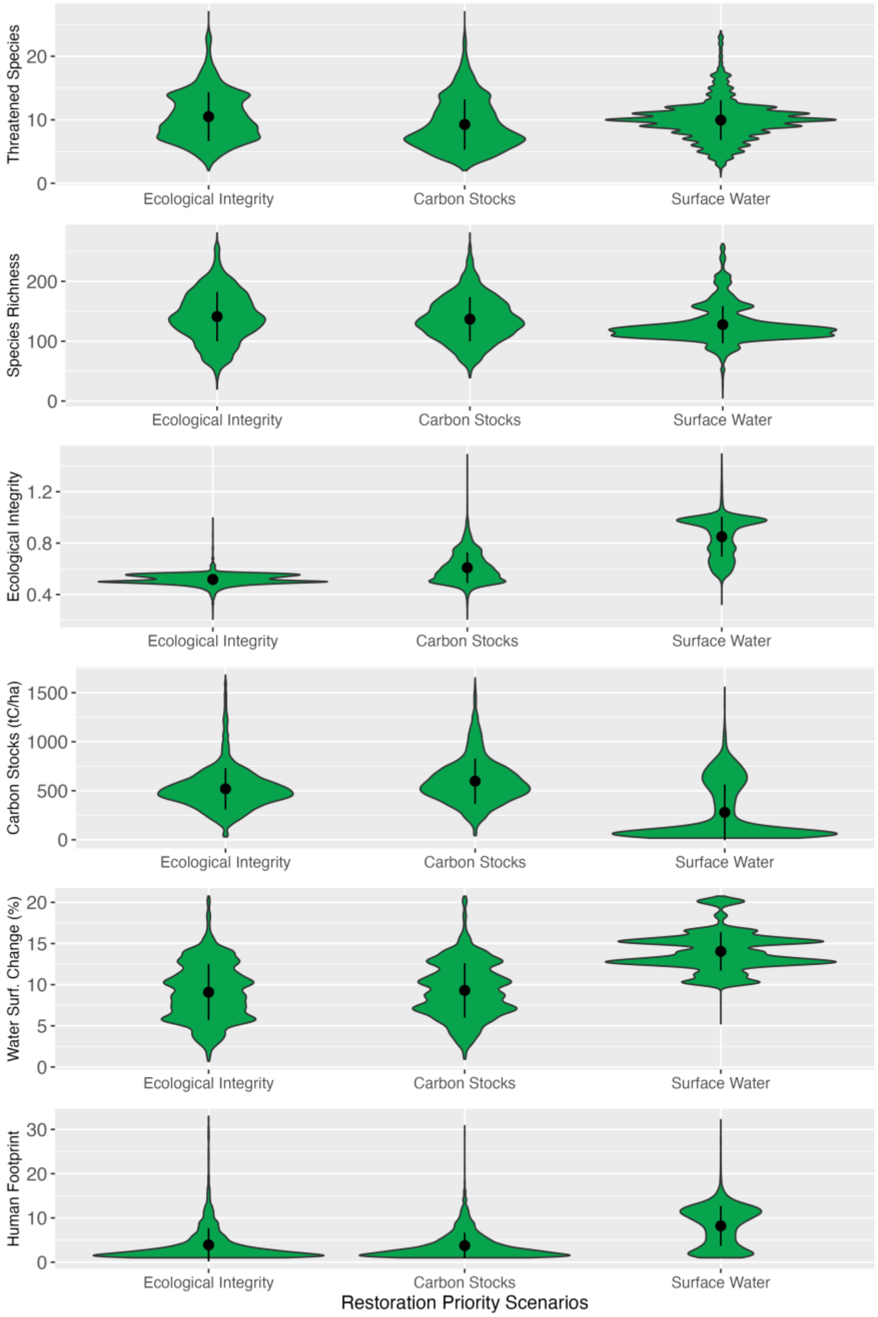


**Figure S11. The distribution of threatened species, species richness, ecological integrity, carbon stocks, absolute surface water change, and human footprint in Restoration Priority Scenarios.** The restoration scenarios aim to maximize the occurrence of threatened species while prioritizing the loss of ecological integrity, carbon stocks, and water surface stability. The points and bars refer to the mean and standard deviation.
